# Using ARCADE (ARChaeplastida Annotation DatabasE) to understand the evolution of genome size in land plants

**DOI:** 10.1101/2022.09.06.506765

**Authors:** Alison Pelri Albuquerque Menezes, João Victor dos Anjos Almeida, Luiz-Eduardo Del-Bem, Francisco Pereira Lobo

**Affiliations:** Departamento de Genética, Ecologia e Evolução, Universidade Federal de Minas Gerais, Belo Horizonte, Minas Gerais, Brazil; Universidade Estadual Paulista Júlio de Mesquita Filho, Jaboticabal, São Paulo; Departamento de Botânica, Universidade Federal de Minas Gerais, Belo Horizonte, Minas Gerais, Brazil

**Keywords:** genome size, C-value, genome evoluttion, genome annotation, comparative phylogenetic methods

## Abstract

The abundance of plant genomic information caused by the decrease of sequencing costs contrasts with the lack of databases that integrate genome annotation, taxonomy and phenotypes to produce statistically sound, biologically meaningful knowledge. Here we present ARCADE (ARChaeplastida Annotation DatabasE), a database of 171 high-quality archaeplastidian non-redundant proteomes gathered from six primary genomic databases, together with proteome quality metrics anda growing number of associated metadata. As a case study to demonstrate the usefulness of ARCADE, we used it to investigate the expansion and contraction of protein domains associated with the evolution of genome size (hereafter GS). GS varies greatly among land plants and the synthesis of large genomes can be costly to cells. Although GS has been studied extensively for decades, the molecular mechanisms involved in the adaptations of plants to the increase in GS are still poorly understood. We used the annotation and phylogenetic information available in ARCADE, together with estimated GS values available for 83 land plant species, to search for associations between the abundance of protein domain families in these species and GS variation through phylogenetic-aware methods. Additionally, we estimated the GS for the ancestral nodes of the extant land plant species. GS seems to be decreasing along the course of evolution, except for a few branches that might have undergone independent GS increases. We found 7 Pfam correlated with the variation in GS in land plants, mainly related to nucleotide metabolism, DNA repair and genome organization. We found larger genomes to have a greater frequency of the Histone 2A superfamily, responsible for diverse functions, including the nucleosome formation and silencing of transposable elements. These molecular functions we found correlated to GS variation suggests they may be associated with preserving genome stability in larger genomes, and might indicate the evolution of mechanisms to cope with the variation in GS in land plants. ARCADE is available at https://bit.ly/ARCADE_OSF.

## 1 INTRODUCTION

Comparative genomics has extensively contributed to elucidate genotype-phenotype associations and understand evolutionary processes at the genome level (Yang et al. 2019). A major conceptual data structure needed for any comparative genomics study is a standardized set of genomic elements (e.g. all protein-coding genes for all species under analysis), each of them annotated to a common dictionary of biologically meaningful annotation terms (e.g. groups of homologous regions shared across gene sets) (Tello-Ruiz et al. 2020). Furthermore, as cellular species are all descendant from the last universal common ancestor and, consequently, are non-independent from a statistical point of view, common association statistics are not suitable to infer genotype-phenotype associations across species (Nagy et al. 2020). For that, a tree-like structure describing the relationships among them is of uttermost importance for model generation, as it allows several phylogeny-aware methods to be applied when searching for genotype-phenotype associations (Adams and Collyer 2017).

Most Archaeplastida members are oxygenic photosynthetic eukaryotes comprising the red algae (Rhodophyta), green algae (Chlorophyta and Charophyta), land plants (Embryophyta), and the freshwater microscopic algae Glaucophytes. Contemporary research in evolutionary plant genomics has been highly impacted by the massive decrease in DNA sequencing costs. The availability of genomic information from early-branching plant lineages, together with comparative genomics analyses, provides the molecular comprehension of major evolutionary phenotypic novelties in this group, such as the vascular system and the flower (Chanderbali et al. 2016; Blázquez, Nelson, and Weijers 2020). These phenotypes are but the tip of the iceberg of a much larger number of quantitative and qualitative phenotypes readily available for comparative analyses across plants (Kattge et al. 2020).

The abundance of plant genomic data contrasts with the challenges associated with the gathering of high-quality, standardized genomic data needed for comparative genomics studies. Several aspects contribute to this paradox. The first and widely known issue is the excess of annotation information available for genes of model organisms when compared with non-model species (Haynes, Tomczak, and Khatri 2018). Another source of bias is the huge variation observed in the quality of genome assemblies due to technical issues, such as distinct sequencing technologies and assembly algorithms, but also caused by true biological facts, such as genome size variation caused by the increase of repetitive elements, domestication events, and lineage-specific whole-genome duplications (Marks et al. 2021). Together, these phenomena can significantly bias downstream comparative genomic analyses.

Not surprisingly, several specialized databases provide high-quality genomic and annotation data for distinct groups of plants. However, these are heavily focused on a few data-rich model organisms comprising mostly angiosperm species of commercial or scientific interest (Marks et al. 2021). This fact limits their usage to provide the genomic and phylogenetic data to survey the evolution of key phenotypic traits in Archaeplastida.

Here we present the ARChaeplastida Annotation DatabasE (ARCADE), a database of high-quality, non-redundant annotated proteomes for 171 Archaeplastida species, together with the available phylogenetic metadata. ARCADE was gathered from six major genome databases and annotated through a common pipeline to provide a rich set of homologous regions as defined by InterProScan, together with Gene Ontology (GO) and pathway annotation, when available (Jones et al. 2014). We also provide phylogenetic information for 142 species as an ultrametric species tree. The integration of phylogenetic and annotation data as provided by ARCADE allows the development of phylogeny-aware models needed for properly comparing species data. Furthermore, instead of focusing on providing user-friendly access to several layers of metadata for a few model organisms and economically relevant angiosperms, as is the case for virtually all plant annotation resources available, we actively searched for high-quality genomes from all known Archaeplastida taxa with an emphasis on underrepresented, early branching lineages, together with their annotation and phylogenetic metadata, when available. This data is available as defined tabular text files and other file formats commonly used in bioinformatics pipelines, therefore providing an annotation and evolutionary scaffold to perform comparative analysis of plants at several taxonomic levels (Nagy et al. 2020).

To demonstrate how our database provides readily available, biologically meaningful knowledge to understand the evolution of complex traits, we used ARCADE to search for homologous regions in protein-coding genes associated with the variation of genome size (or C-value) in land plants, whose variation has been largely studied and yet is one of the greatest mysteries of genomics and evolutionary biology. The associations of genome size variation and several plant traits have been repeatedly addressed. However, the molecular mechanisms involved in the adaptations of plants to the increase in genome size are still poorly understood. Thus, we aim at using this case study to demonstrate ARCADE’s usefulness by investigating which protein domain famlies are correlated with the increase in genome size in Embryophyta (land plants). There might be molecular mechanisms that allow genome size variation in nature, considering that the increase in genome size might also impose physiological pressures on their organisms (Knight, Molinari, and Petrov 2005). Through analysis of comparative genomics on 83 species of land plants, we found that protein domains associated with key components of DNA biology, such as DNA repair, nucleotide metabolism and genome maintenance are enriched in plants with larger genomes when compared to the smaller ones. These results might show the way to mechanisms through which plants cope with the impact of genome increase, and comprise a compelling case study of the usefulness of our database.

## 2 METHODS

### 2.1 Building ARCADE

#### 2.1.1 Survey for Archaeplastida complete genomes

We screened six primary genomic databases for plant genomes: 1) National Center for Biotechnology Information (NCBI) databases – RefSeq (O’Leary et al. 2015) and GenBank (Sayers et al. 2019); 2) Phytozome (Goodstein et al. 2011); 3) Gymno PLAZA (Proost et al. 2014); 4) FernBase (F.-W. Li et al. 2018); 5) CNGB (X. Wang et al. 2010); 6) and the DRYAD repository (J. Zhang et al. 2020). We aimed at building a comprehensive and phylogenetic diverse genomic dataset for Archaeplastida species, with emphasis on early-branching lineages. Specifically, we downloaded the assembled genomic data for all Embryophyta species available in the NCBI databases and complemented our dataset with the predicted proteomes belonging to relevant and underrepresented lineages with data from the other public databases.

#### 2.1.2 Obtention of high-quality, non-redundant annotated proteomes and phylogenetic data

To reduce the proteome redundancy generated by the distinct number of isoforms of the same gene and avoid a possible bias towards model organisms, we applied two different proteome summarization protocols to our dataset. The first protocol, used on NCBI data (*in-house* pipeline), selects the longest protein sequence of each protein-coding locus based on the “locus_tag” or “gene_id” fields. However, fasta proteomes from other databases do not contain these fields in their metadata, and could not be submitted to our pipeline. To reduce redundancy of those proteomes we used the CD-HIT software (W. Li and Godzik 2006; Fu et al. 2012) with the threshold set to 1; this way we expected to keep only the longest protein isoform to represent each genomic locus. We evaluated the assembling quality of each proteome using the BUSCO software (Manni et al. 2021) to assess their gene completeness based on the set of 255 Viridiplantae BUSCOs (odb10). We kept in our dataset proteomes with completeness higher than 70% and rates lower than 20% for duplicated and fragmented genes, using the results obtained for *Arabidopsis thaliana* as a proxy for the lower-bound cutoffs for quality (Supplementary Table S1).

We used InterProScan 5 (Jones et al. 2014) to perform a *de novo* annotation of the 171 non-redundant proteomes that fulfill the previous BUSCO cutoffs (from now on referred as high-quality non-redundant proteomes -NRPs). Specifically, all NRPs were annotated to 13 distinct databases that are integrated into InterProScan that provide homology and functional annotation information for all proteomes, if available.

The ARCADE database comprises a tabular file containing curated metadata for all 171 species, such as species binomial names, NCBI TaxonIDs, database of origin and proteome quality metrics, among others. We also provide annotation information, including raw output files from InterProScan for each species, as well as parsed data for all distinct annotation databases members of InterProScan. Finally, we also gathered a newick file containing a species tree for 142 species gathered from the TimeTree web tool (Kumar et al. 2022). The initial tree produced by this tool was enriched by including species from closely related taxa as placeholders for the original species. Example: if one species from an order could not be placed on the tree and 1) it is the only species from this order present in ARCADE and 2) another species from the same order is available, we replaced the original species by the one present in TimeTree to include the original species in the tree.

### 2.2 Case study – evolution of genome size in Embryophyta

#### 2.2.1 Computation environment

The analysis done in this project was executed in a Dell server with 2 processors Intel Xeon E5-4610 v2 2.3GHz totalizing 64 threads; 128GB of RAM and operational system CentOS Linux release 7.5.1804, with support to the programming languages PERL5 v. 7.5 and R v. 3.0.0.

#### 2.2.2 Data Collection and Functional Annotation

As ARCADE contains species’ scientific names as its primary key to integrate distinct data types, it is trivial to gather information from other databases and integrate with the genomic annotations we provide. To investigate the evolution of genome size in land plants, we obtained experimentally measured 1C-value data for all the studied species in the Plant C-values Database (Pellicer and Ilia J. Leitch 2020) and in the GoaT database (Challis et al. 2017) (Supplementary Table S1). Here we are using 1C-value and genome size convertibly, although the genome size would equal to 2C-value divided by ploidy level (Ilia J. Leitch, Chase, and Bennett 1998). Then, we converted the 1C-value from picogram (pg) to megabase pairs (Mb), where 1 pg equals 980 Mb and log10-transformed those values. To evaluate the distribution of GS values in our sample we performed the Shapiro-Wilk test (SHAPIRO and WILK 1965). In the end, we gathered genotypic, phylogenetic, and genotypic data for 83 species of Embryophyta (Supplementary Table S1, Fig. 1) to perform our phylogenetic comparative analysis.

**Figure 1:**
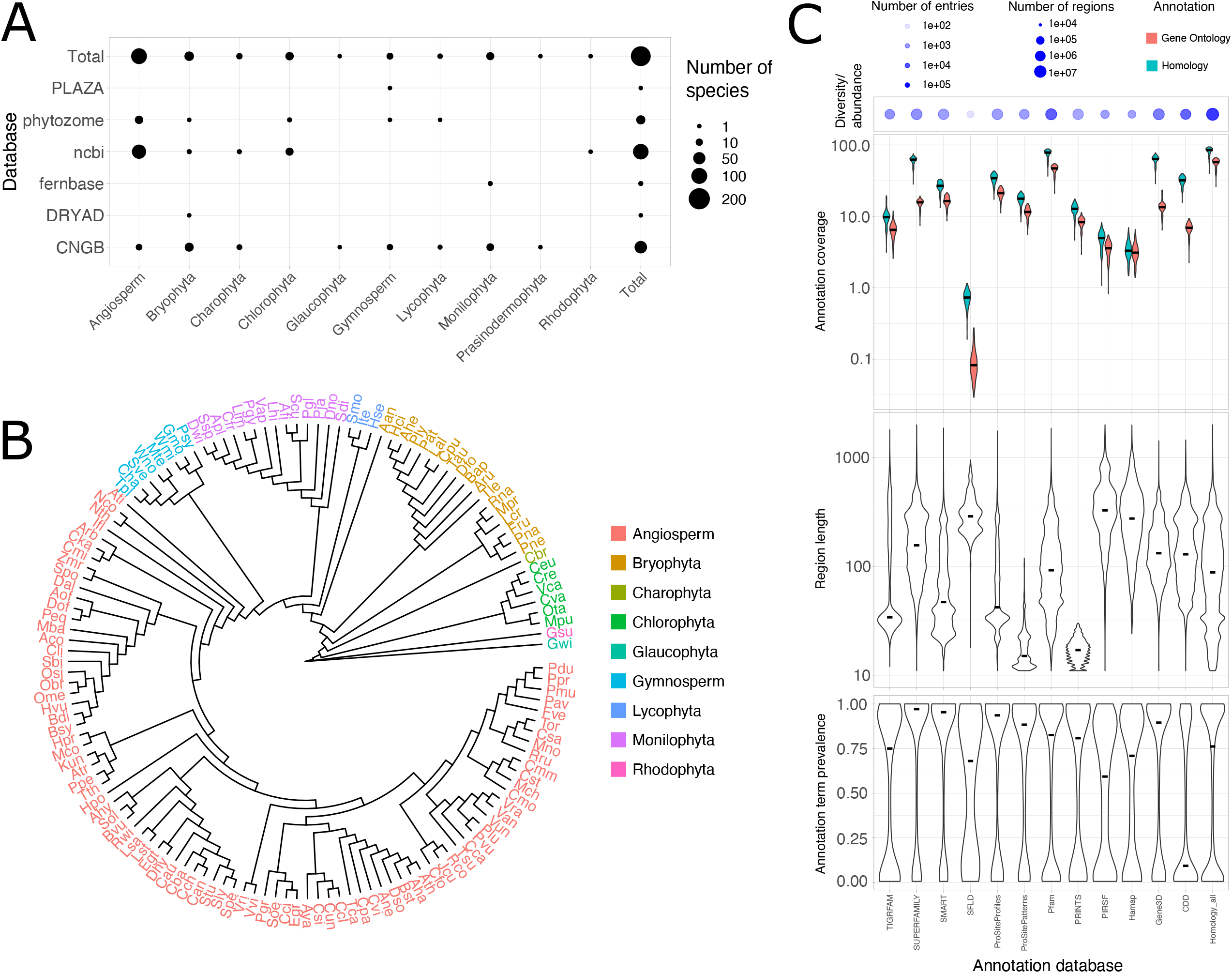
Description of ARCADE database. A) Relative species richness and taxonomic diversity of the plant genome databases used to build ARCADE. B) Species tree for 142 species found in ARCADE (extracted from the TimeTree web tool (Kumar et al. 2022) as described in “Methods”. Species abbreviations are as follows: *Acer yangbiense* (Aya), *Akebia trifoliata* (Atr), *Amaranthus hypochondriacus* (Ahy), *Amborella trichopoda* (Ati), *Ananas comosus* (Aco), *Andreaea rupestris* (Aru), *Anthoceros angustus* (Aan), *Arabidopsis halleri* (Aha), *Arabidopsis thaliana* (Ath), *Arabis nemorensis* (Ane), *Ascarina rubricaulis* (Arb), *Asparagus officinalis* (Aof), *Asplenium platyneuron* (Apl), *Aulacomnium heterostichum* (Ahe), *Azolla filiculoides* (Afi), *Beta vulgaris subsp. vulgaris* (Bvu), *Boechera stricta* (Bst), *Brachypodium distachyon* (Bdi), *Brachypodium sylvaticum* (Bsy), *Buxbaumia aphylla* (Bap), *Cajanus cajan* (Cca), *Cannabis sativa* (Csa), *Capsicum annuum* (Can), *Capsicum baccatum* (Cba), *Capsicum chinense* (Cch), *Carex littledalei* (Cli), *Carica papaya* (Cpa), *Castanea mollissima* (Cmo), *Cephalotaxus harringtonia* (Cha), *Cephalotus follicularis* (Cfo), *Ceratodon purpureus* (Cpu), *Chara braunii* (Cbr), *Chlamy-domonas eustigma* (Ceu), *Chlamydomonas reinhardtii* (Cre), *Chlorella variabilis* (Cva), *Cinnamomum kanehirae* (Cka), *Cinnamomum micranthum* (Cmi), *Citrus clementina* (Ccl), *Citrus sinensis* (Csi), *Citrus unshiu* (Cun), *Cleome violacea* (Cvi), *Coleochaete irregularis* (Cir), *Corymbia citriodora* (Cci), *Cucumis melo* (Cme), *Cucumis sativus* (Cst), *Cuscuta australis* (Cau), *Cystopteris fragilis* (Cfr), *Danaea nodosa* (Dno), *Dendrobium officinale* (Dof), *Descurainia sophioides* (Dso), *Dioscorea alata* (Dal), *Diphyscium foliosum* (Dfo), *Diplazium wichurae* (Dwi), *Dorcoceras hygrometricum* (Dhy), *Erythranthe guttata* (Egt), *Eucalyptus grandis* (Egr), *Fragaria vesca subsp. vesca* (Fve), *Frullania spp*. (Fru), *Galdieria sulphuraria* (Gsu), *Gloeochaete wittrockiana* (Gwi), *Gnetum montanum* (Gmo), *Hakea prostrata* (Hpo), *Hedwigia ciliata* (Hci), *Hordeum vulgare* (Hvu), *Huperzia selago* (Hse), *Hypecoum procumbens* (Hpr), *Illicium floridanum* (Ifl), *Isoetes tegetiformans* (Ite), *Jatropha curcas* (Jcu), *Kingdonia uniflora* (Kun), *Lactuca saligna* (Lsa), *Lactuca sativa* (Lst), *Leucobryum albidum* (Lal), *Leucostegia immersa* (Lim), *Lonchitis hirsuta* (Lhi), *Lunularia cruciata* (Lcr), *Macleaya cordata* (Mco), *Marchantia polymorpha* (Mpo), *Microcachrys tetragona* (Mte), *Micromonas pusilla CCMP1545* (Mpu), *Momordica charantia* (Mch), *Morus notabilis* (Mno), *Musa balbisiana* (Mba), *Nymphaea colorata* (Nco), *Nymphaea thermarum* (Nth), *Oryza brachyantha* (Obr), *Oryza meyeriana var. granulata* (Ome), *Oryza sativa* (Osa), *Ostreococcus tauri* (Ota), *Pellia neesiana* (Pne), *Phalaenopsis equestris* (Peq), *Phaseolus lunatus* (Plu), *Phaseolus vulgaris* (Pvu), *Pilularia globulifera* (Pgl), *Pinus sylvestris* (Psy), *Plenasium javanicum* (Pja), *Podophyllum peltatum* (Ppe), *Polypodium glycyrrhiza* (Pgy), *Porella navicularis* (Pna), *Prunus avium* (Pav), *Prunus dulcis* (Pdu), *Prunus mume* (Pmu), *Prunus persica* (Ppr), *Pseudanomodon attenuatus* (Pat), *Pulvigera lyellii* (Ply), *Punica granatum* (Pgr), *Rhamnella rubrinervis* (Rru), *Rhododendron williamsianum* (Rwi), *Ricciocarpos natans* (Rna), *Ricinus communis* (Rco), *Salvinia cucullata* (Scu), *Sceptridium dissectum* (Sdi), *Sciadopitys verticillata* (Sve), *Selaginella moellendorffii* (Smo), *Solanum lycopersicum* (Sly), *Solanum pennellii* (Spe), *Solanum tuberosum* (Stu), *Sorghum bicolor* (Sbi), *Spinacia oleracea* (Sol), *Spirodela polyrhiza* (Spo), *Struthiopteris spicant* (Ssp), *Syzygium oleosum* (Soe), *Takakia lepidozioides* (Tle), *Thalictrum thalictroides* (Tth), *Theobroma cacao* (Tca), *Thuja plicata* (Tpl), *Timmia austriaca* (Tau), *Trema orientale* (Tor), *Trifolium subterraneum* (Tsu), *Vigna angularis* (Van), *Vigna radiata var. radiata* (Vra), *Vigna unguiculata* (Vun), *Vitis riparia* (Vri), *Vitis vinifera* (Vvi), *Vittaria appalachiana* (Vap), *Volvox carteri f. nagariensis* (Vca), *Welwitschia mirabilis* (Wmi), *Wollemia nobilis* (Wno), and *Zostera marina* (Zmr). C) Characterization of the 12 InterPro databases used to annotate ARCADE sequence data. Plots represent information as follows: Diversity/abundance – the number of entries

#### 2.2.3 Comparative analysis

We studied the evolution of genome size in land plants in two steps. First, we estimated the ancestral genome size and their respective 95% confidence interval (CI) for all the internal nodes of the obtained phylogeny based on the genome size of 86 extant species of land plants. That was made by the method of maximum likelihood implemented in the “*fastAnc*” function of the *phytools* R package (Revell 2012). Then we mapped the estimated ancestral states (*fastAnc* function) on the phylogeny obtained from the Time Tree of Life database (Kumar et al. 2022) using ggplot2 (Sievert 2016).

Second, we investigated protein domains’ abundance and frequency correlated with the increase in genome size in land plants. To do so, we used CALANGO (Hongo et al. 2021), which integrates protein domain annotation (both frequency and absolute abundance in each genome), genotypic (genome size), and phylogenetic data. To avoid bias from the phylogenetic interdependencies within the data, the CALANGO applies the method of Phylogenetically Independent Contrasts (PIC) (Felsenstein 1985) using the pruned species phylogenetic tree distributed with ARCADE.

To summarize results and facilitate data interpretation, the Pfam domains obtained from the results of the CALANGO analysis were used to find, in the InterProScan annotation files, the *Arabidopsis thaliana* genes containing them. Then we searched for information about those genes using the ThaleMine (Krishnakumar et al. 2014) (Pasha et al. 2020) data mining tool to gather relevant functional information.

## 3 RESULTS AND DISCUSSION

### 3.1 The ARCADE database: a brief overview

We created ARCADE as a resource to organize and integrate high-quality non-redundant predicted proteomes from Archaeplastida species, together with their respective phylogenetic metadata and genomic annotations, when available. For that, we surveyed six major genome databases that host plant genomes and, starting from 1,381 species, we gathered 171 high-quality, non-redundant proteomes of Archaeoplastida organisms from the following ten major groups (number of species between parenthesis): Angiosperm (98), Gymnosperm (8), Bryophyta (24), Monilophyta (14), Charophyta (6), Chlorophyta (16), Glaucophyta (1), Lycophyta (3), Prasinodermophyta (1), Rhodophyta (1) (Figure 1A; see also section “Comparison of ARCADE and other plant genome databases” for a deeper discussion on individual database contribution for the final list of species available in ARCADE). We could confidently produce a species tree for 142 species found in ARCADE (84.2%) that represent all major taxa but Prasinodermophyta, therefore providing a taxonomically diverse phylogenetic scaffold for phylogeny-aware studies, as we demonstrate in our case study (Figure 1B).

Model organisms are data-rich species and expected to have more genomic information available, including the number of known isoforms, and these may be a significant source of bias (Chen et al. 2014), and non-redundant proteomes are a mainstream source of unbiased genomic components for comparative genomic studies (Vogel and Chothia 2006). We used locus information available at NCBI to generate non-redundant proteomes for all species. However, as genomes from all other databases do not provide isoform-level information, we proceed by using CD-HIT to remove redundancy from the predicted proteomes and evaluated whether this strategy also introduced any undesirable downstream bias. For that we compared the non-redundant proteomes of four major groups summarized by both methods (in house and CD-HIT): angiosperms, bryophytes, charophytes and chlorophytes. Specifically, we evaluated these groups for the number of protein sequences found in each proteome, together with BUSCO metrics for proteome completeness (Fig. S1). We found CD-HIT summarization to produce, on average, smaller proteomes for three out of the four groups (angiosperms, bryophytes and chlorophytes), with an opposite trend observed for charophytes (Fig. S2). The BUSCO results for completeness and single-copy BUSCOs displayed a similar trend (Fig. S3-4).

The number of duplicated BUSCOs, which are a widespread proxy to detect genome assemblies with duplications introduced by genome assembly issues, and is also expected to be higher in proteomes with more than one isoform per locus, showed similar distributions for all four groups but bryophytes, where CD-HIT summarization strategy appears to produce a higher number of duplicated genes (Fig. S5). Up to this point, one could conclude that the CD-HIT may be introducing bias in at least the non-redundant proteomes from bryophyte species. However, the analysis of fragmented and missing BUSCOs, which are likely not to be influenced by any summarization protocol, found that proteomes summarized by CD-HIT more fragmented and have more missing BUSCOs than their *in-house* counterparts, especially for the bryophytes (Supplementary Figures S6-7). Taken together, we conclude that the CD-HIT summarization strategy does not introduce any significant bias during the production of non-redundant proteomes. The differences observed are likely due to other factors, such as true natural variation or genome assembly issues. Although databases other than NCBI were very important to increase diversity in our dataset, our results for BUSCO values suggest that in general, their assembly quality is lower than the NCBI ones.

The annotation of all 171 high-quality non-redundant proteomes using the 12 databases as provided by InterProScan predicted a total of 23,173,794 homologous regions within non-redundant proteomes annotated by one of the 23,514 distinct annotation terms, defined as each distinct homologous region from a specific annotation database. We found these databases to provide highly variable sets of annotation terms for sequence length, taxonomic prevalence and functional diversity (Figure 1C; group “Homology_all” represents all entries from all databases). There is a considerable variation in annotation term abundance (regions annotated by an annotation term) and diversity (number of distinct annotation terms) contributed by each database. Two databases (Pfam and CDD) have approximately 25% and 50% of the total annotation abundance and diversity, respectively (group Figure 1C, “Diversity/abundance” plot).

We also found distinct databases to have large differences in the annotation coverage of non-redundant proteomes, defined as the fraction of non-redundant proteomes annotated by at least one annotation term. Three databases (Pfam, SUPERFAMILY and GENE3D) provide the broadest values of proteome annotations. Importantly, a considerable fraction of the proteomes found in ARCADE are annotated by at least one annotation term (Figure 1C, “Annotation coverage” plot, “Homology_all” group). In contrast with all other annotation databases, Pfam also contributes with the largest fraction of Gene Ontology (GO) annotation. The annotation databases are also highly variable in terms of the length of each homologous region, ranging from databases enriched in short sequences (e.g. PRINTS, ProSitePatterns) to databases with median length homologous regions around 300 amino acid residues (e.g. PIRSF).

Finally, we evaluated annotation term prevalence across proteomes, defined as the fraction of proteomes where an annotation term was found (Figure 1C, “Annotation term prevalence” plot). We observed databases to possess a bimodal distribution with annotation terms that have either low prevalence, being observed in few proteomes and corresponding to lineage-specific homologous regions, to annotation terms highly prevalent, comprising homologous regions found in the majority of the species under analysis. The vast majority of databases are enriched in highly prevalent homologous regions, with CDD being the only database enriched in homologous genes with low prevalence across species. Taken together, we conclude that ARCADE represents a considerable effort towards providing standardized and high-quality phylogenetic and annotation data, where individual annotation databases provide a highly variable set of homologous regions in terms of abundance, diversity, phylogenetic distribution, annotation coverage and functional characterization.

### 3.2 Comparison of ARCADE and other plant genome databases

The genome databases that host data for plant species, such as the ones we used to mine data from, are developed for scientists with little or no computational background. As so, they provide rich graphical user interfaces to allow data access and analysis through genome browsers and the integration of several layers of genomic metadata. However, these databases mostly host data from specific lineages, or have a phylogenetically limited and biased dataset focused on model species or of agricultural interest (Ma et al. 2022). ARCADE, in contrast, was specifically developed to provide a taxonomically diverse selection of high-quality, non-redundant predicted proteomes, together with its phylogenetic and annotation data, if available. Our target audience is the community of plant computational biologists that may take advantage of having a taxonomically diverse set of species and their fundamental data types needed for the study of complex genotype-phenotype associations. More importantly, the data provided in ARCADE is not found in any single plant genome database, as we demonstrate below.

NCBI, CNGB, and Phytozome are some of the richest resources in plant genomic data. NCBI databases are arguably the most commonly used resources for genetic and genomic studies in the western world, thus it gathers a very wide range of different types of data, very well curated and organized. CNGB is a large-scale database with an enormous number and diversity of genomes, but it does not appear to be well curated. From thousands of predicted proteomes, only 54 unique species were approved by our quality control. Furthermore, we found BUSCO results to be consistently worst from genomes summarized by CD-HIT, as is the case for all CNGB entries (Supplementary Figures S3-S7).

Even though a single database (NCBI) contributes with the majority of entries of ARCADE (94 spp, 60,0%), no individual genomic database was found to contribute with species for all ten major Archaeplastida groups, with CNGB and NCBI being the most taxonomically rich databases and contributing with 8 and 5 groups, respectively. Furthermore, no single group was found in all genomic databases, with Bryophyta being the most prevalent group (4 out of 6 databases), and with three major groups observed in a single database (Rhodophyta – NCBI, Prasinodermophyta and Glaucophyta – CNGB). Among all the databases we searched, Phytozome is the only one containing functional annotations for their predicted proteomes and the most phylogenetically diverse as well. The databases that bring functional annotation to their protein sequences are more specific, focusing on tRNA (Chan and Lowe 2015) or specific enzyme families (Ekstrom et al. 2014), for example. Phytozome is well-curated and offers comprehensive and uniform annotation for the genomes of its 139 Archaeplastida species, while our datasets encompass 171 plant species. Although Phytozome includes more phylogenetic diverse species, the number of representants of non-flowering plants is insufficient, even lacking species of the major clade of charophytes.

We also used genomes from lineage-specific databases, such as FernBase and Gymno PLAZA, to increase the diversity of ferns and gymnosperms in our dataset. Only *Anthoceros angustus* did not come from a genomic database; this proteome was obtained from the data repository DRYAD where it was uploaded by the research group responsible for sequencing its genome. Taken together, we consider that our effort to obtain a diverse set of high-quality and annotated non-redundant proteomes for Archaeplastida was successful, as these would not be obtainable from any single database we evaluated.

### 3.3 Case study: using ARCADE to study the evolution of genome size in land plants

The genome size varies about 2,400-fold in land plants, ranging from 61 Mb in *Genlisea tuberosa* (the smallest genome known yet) (Fleischmann et al. 2014) to 152,000 Mb in *Paris japonica* (Pellicer, Fay, and Ilia J. Leitch 2010). Since genome sizes are readily available since long before the beginning of genomic studies, variations on this trait have been largely evaluated for associations (Bennet and Ilia J. Leitch 2005). Previous studies have found correlations between genome size and many plants’ phenotypic and ecological traits, such as cell and seed size, habitat, and distribution (Bennett 1987; Knight, Molinari, and Petrov 2005; Beaulieu et al. 2007; Kang et al. 2014).

In land plants, large genomes emerged mainly due to repeated events of polyploidy and the imbalance between the amplification and removal of transposable elements from DNA (Pellicer, Hidalgo, et al. 2018; Petrov 2001). Genome size imposes restrictions on compatible life strategies and ecological options, as it will affect plant development and fitness (Knight, Molinari, and Petrov 2005; A. R. Leitch and I. J. Leitch 2012). Due to the costs of maintaining their genome (e.g. nutrient and water supply), plants with larger genomes will be limited to more stable environments, where selective pressures are more relaxed (Knight, Molinari, and Petrov 2005; Veselý, Bureš, and Šmarda 2013; Hidalgo et al. 2017). Many authors have proposed different evolutionary models that place genome size as a trait that responds to selective pressures, adaptively or not, or as an evolutionary neutral trait. Therefore, it is fair to assume that genome size variation holds biological importance (Petrov 2002; Vinogradov 2003).

To demonstrate how ARCADE provides the phylogenetic and annotation scaffold needed to survey the evolution of complex traits, we used our dataset to survey the influence of genome size (C-value) variation in land plants and the abundance of homologous genes as defined by the Pfam annotation (Fig. S2 contains the workflow for this analysis). We started by gathering the C-values for 86 species of land plants found in ARCADE. At this point, it is worth noting that five out of the six genome databases used to generate ARCADE contributed with species for this case study, therefore demonstrating the unfeasibility of using any individual database to reproduce our study case study and, consequently, the usefulness of ARCADE.

#### 3.3.1 Integrating phylogenetic and trait data to estimate ancestral of genome size

Shapiro-Wilk test showed that the distribution of C-values is not normal, even after log10-transformation (W = 0.89674, p < 0.0001), with a long-tail towards larger genomes (Fig. 2). The range of C-values varies from 86.24 Mb (for the lycophyte *S. moellendorffii*) to 24,792.30 Mb (for gymnosperm *Cephalotaxus harringtonia*). The genome size average in our dataset is 2,394.33 Mb, but 67 species (of 86 in total) have genomes smaller than the average, dragging the median down to 618.63 Mb. Most plants with large genomes are gymnosperms (14,992.32 Mb on average) or monilophytes (4,258.59 Mb on average) (Figure 2). But there are also representatives of the plants with large genomes within angiosperms (i.e. *Hordeum vulgare* with 5,379 Mb) and lycophytes (i.e. *Huperzia selago* with 4,938.9 Mb). We used current C-values, together with the species tree provided by ARCADE, to estimate the genome size for the ancestral nodes of the land plant phylogeny (Figure 2A contains the estimations; Figures 2B-C contains the actual values for species and major groups, respectively).

**Figure 2:**
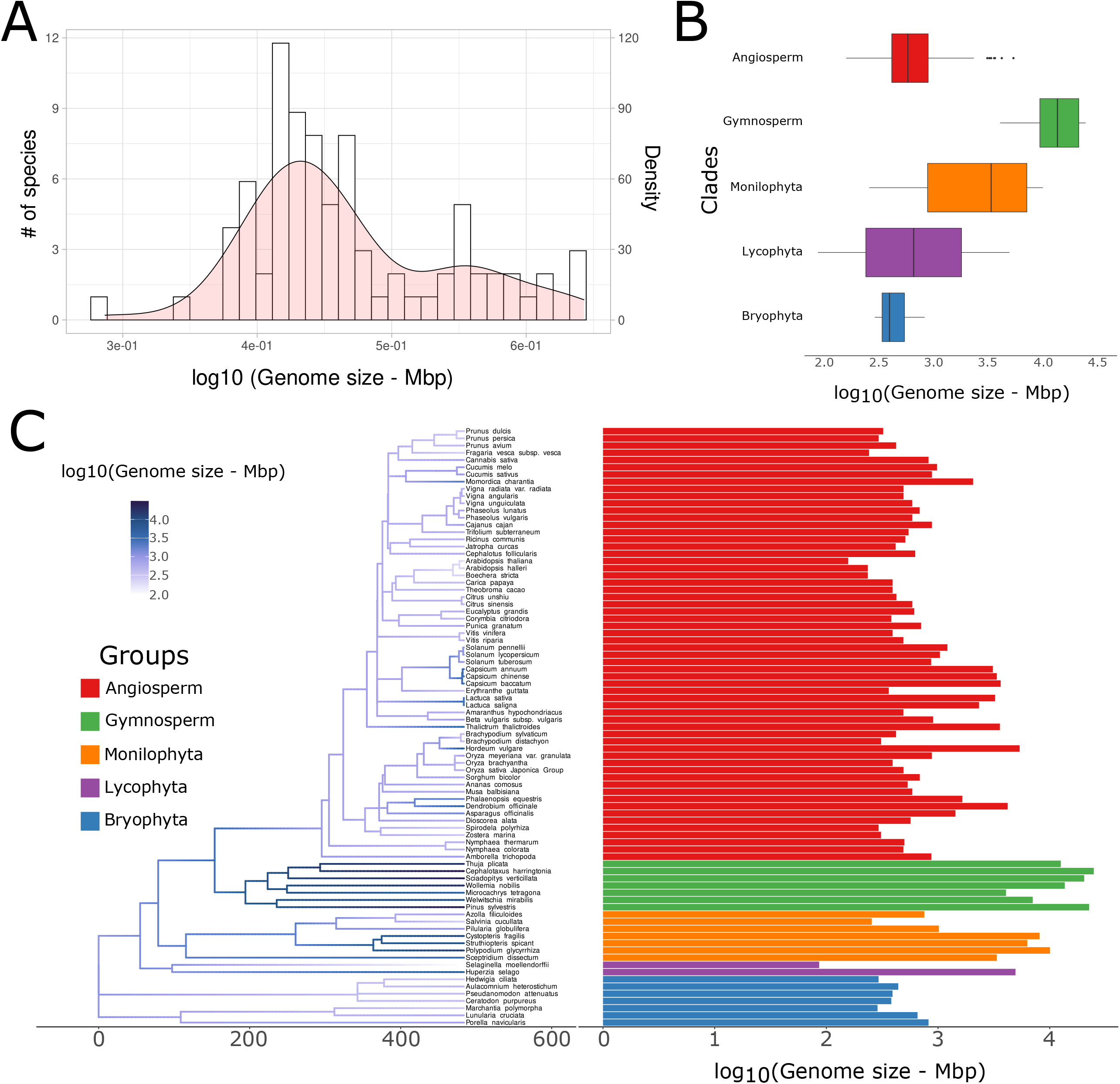
Evolution of genome size in land plants. A) Overall distribution of genome size in land plants. B) Variation of genome size in groups of land plants. C) Estimation of ancestral genome size values.

The estimated ancestral genome size of the land plants was 1088.98 Mb (95% CI: 212.93 Mb to 5,569.38 Mb), smaller than the average genome size of the extant plants. From that point in evolution, it shows bidirectional evolution. The GS in most clades are decreasing, but we can notice that in few clades GS is increasing independently along the phylogeny of land plants, as previously observed (Bennett 1987; Wendel et al. 2002).

The last common ancestor of all spermatophytes is estimated to have a genome of 2916.91 Mb (95% CI: 789.24 Mb to 10780.43 Mb), following an increase in genome size in Gymnosperms, with the largest genomes, as mentioned before and a last common ancestor with a genome size of 5265.78 Mb (95% CI: 1535 Mb to 18064.16 Mb). Then, there is a pattern of decrease in genome size within the Angiosperm clade, with a genome of 958.67 Mb (95% CI: 324.80 Mb to 2829.53 Mb) in their common ancestor and an average of 2457.63 Mb among the extant species of angiosperms. However, we can notice that a few lineages of angiosperms went through an increase in genome size, such as the ancestral state of the common ancestor of Asteraceae, Orchidaceae, and Solanaceae, for which the estimated genome size of their common ancestor was 2742.47 Mb (95% CI: 2575.76 Mb to 2919.96 Mb), 1675.07 Mb (95% CI: 686.5 Mb to 4086.64), and 1607.76 Mb (95% CI: 944.53 Mb to 2736.68), respectively. On the other hand, in Mosses, we observe an average of 376.53 Mb, and the common ancestor genome size estimative is 417.36 Mb (95% CI: 129.85 Mb to 1341.49 Mb). Similarly, in Liverworts the average genome size is 587.97 Mb and the estimative for the genome size of the last common ancestor is 842.49 Mb (95% CI: 131.84 Mb to 5383.54 Mb). We can notice close species from other lineages with both increasing and decreasing genome size. There are two representatives in the Lycophyta branch, *S. moellendorffii* with the smallest genome in our dataset, 86.24 Mb, and *H. selago* with 4938.9 Mb of genome size, among the largest ones.

Almost every node and branch tip in the bryophyte clades (mosses and liverworts) present a decrease in genome size, except for the *Aulacomnium heterostichum*, which has a genome of 655.26 Mb and descends from a node estimated to have a genome size of 393.17 Mb. However, both clades maintain their genome sizes very small, as described before for bryophytes (Bennet and Ilia J. Leitch 2005).

Every other clade has examples of branches both increasing and decreasing the genome size, which is consistent with the literature (Bennet and Ilia J. Leitch 2005). Lycophytes and ferns are both very diverse clades concerning genome sizes (D. Wang et al. 2021), thus great cases to observe the variation of genome size within a clade. Although we have a small sample of lycophyte representatives, they interestingly are *Selaginella moellendorffii* (the smallest genome in our dataset) and *H. selago* (among the largest ones in our dataset). On the other hand, in ferns, it is easy to observe the different patterns of genome size evolution, while the Salvinales show a clear decrease in genome size, we can see the genome size of Polypodiales increasing, some of the largest in the dataset.

The estimated genome size for the ancestor of Spermatophyta is 2916.91 Mb and, again, we notice two different pattern behaviors. In gymnosperms, the genomes sizes are uniform within the clade (Puttick, Clark, and Donoghue 2015) and increase along the phylogeny, except for *Welwitschia mirabilis* and *Microcachrys tetragona*. Bennet and Ilia J. Leitch (2005) also demonstrated this genome downsizing in the Gnetales, as the *W. mirabilis*, suggesting a contraction of genome size in the order (Bennet and Ilia J. Leitch 2005). Displaying a different trend from gymnosperms, the angiosperms show a high diversity of genome size, but mostly very small genome sizes, from its common ancestor to most of the nodes and branches within the clades (I. J. Leitch et al. 2005). Although most genomes are very small, few derived clades evolved independently intermediate genomes, specially Solanaceae, Orchidaceae, and Asteraceae (Bennet and Ilia J. Leitch 2005). The patterns of genome size distribution and diversity in angiosperms could explain the success and diversity of angiosperm species, as genome size is known to be correlated (directly or indirectly) to several functional and adaptive traits (Ilia J. Leitch, Chase, and Bennett 1998; Bennet and Ilia J. Leitch 2005; Puttick, Clark, and Donoghue 2015; Carta et al. 2022).

#### 3.3.2 Protein domains associated with genome size in land plants

We used CALANGO, an *in-house* comparative genomics tool that integrates genotypic, phenotypic and annotation data, to build phylogeny-aware linear models and search for phenotype-genotype associations, to investigate possible homologous regions associated with the variation of genome size in land plants (Hongo et al. 2021). Specifically, we used 1) the the log10(C-value) and 2) either the absolute counts or the relative frequencies (the ratio of each Pfam annotation count to the total number Pfam annotation in each proteome) of annotations terms defined by Pfam as data vectors for phenotypes and genotypes, respectively. From a total of 5,721 Pfam IDs that annotate at least one protein-coding gene from one of the 83 genomes of land plants under analysis, we found that 7 of them (0.12%) are significantly correlated with the increase in genome size in land plants (q-value < 0.1) (Table 1, Figure 3). All of them were found in at least 90% of the studied species across the whole phylogeny of land plants (prevalence > 0.9), suggesting these are conserved homologous regions shared across most land plants in this analysis.

**Table 1:**
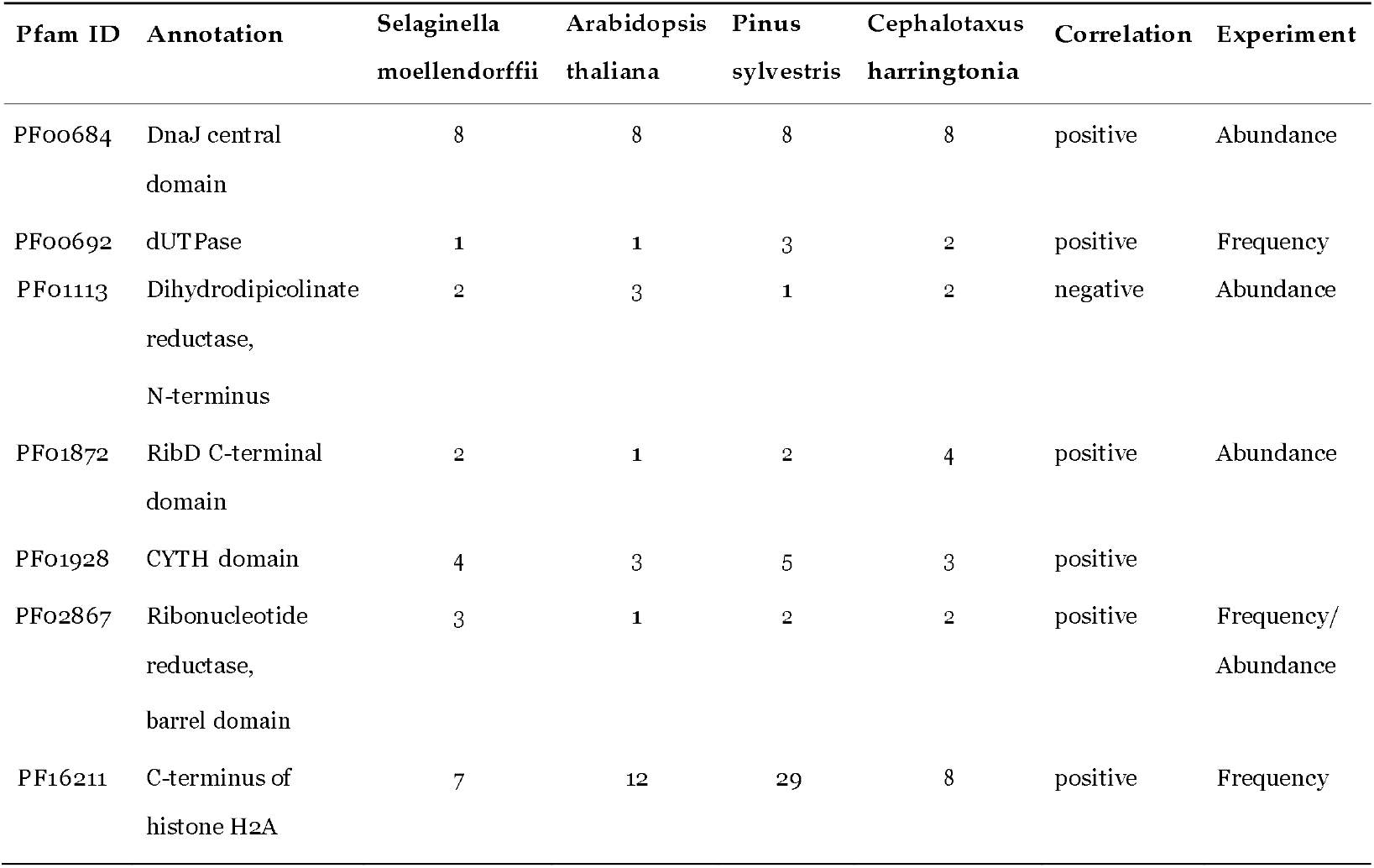
Protein domains found significantly correlated to the genome size evolution in land plants and their respective annotations, number of copies in the two species with shortest (*Selaginella moellendorffii, Arabidopsis thaliana*) and largest (*Pinus sylvestris, Cephalotaxus harringtonia*) genomes in our dataset, the signal of the correlation, and whether the correlation is regarding the Pfam domain’s abundance or frequency.

| Pfam ID | Annotation | Selaginella<br>moellendorffii | Arabidopsis<br>thaliana | Pinus<br>sylvestris | Cephalotaxus<br>harringtonia | Correlation | Experiment |
| --- | --- | --- | --- | --- | --- | --- | --- |
| PFO0684 | DnaJ central domain | 8 | 8 | 8 | 8 | positive | Abundance |
| PFO0692 | dUTPase | 1 | 1 | 3 | 2 | positive | Frequency |
| PFO1113 | Dihydrodipicolinate reductase, N-terminus | 2 | 3 | 1 | 2 | negative | Abundance |
| PFO1872 | RibD C-terminal domain | 2 | 1 | 2 | 4 | positive | Abundance |
| PFO1928 | CYTH domain | 4 | 3 | 5 | 3 | positive |  |
| PFO2867 | Ribonucleotide reductase, barrel domain | 3 | 1 | 2 | 2 | positive | Frequency/<br>Abundance |
| PF16211 | C-terminus of histone H2A | 7 | 12 | 29 | 8 | positive | Frequency |

**Figure 3:**
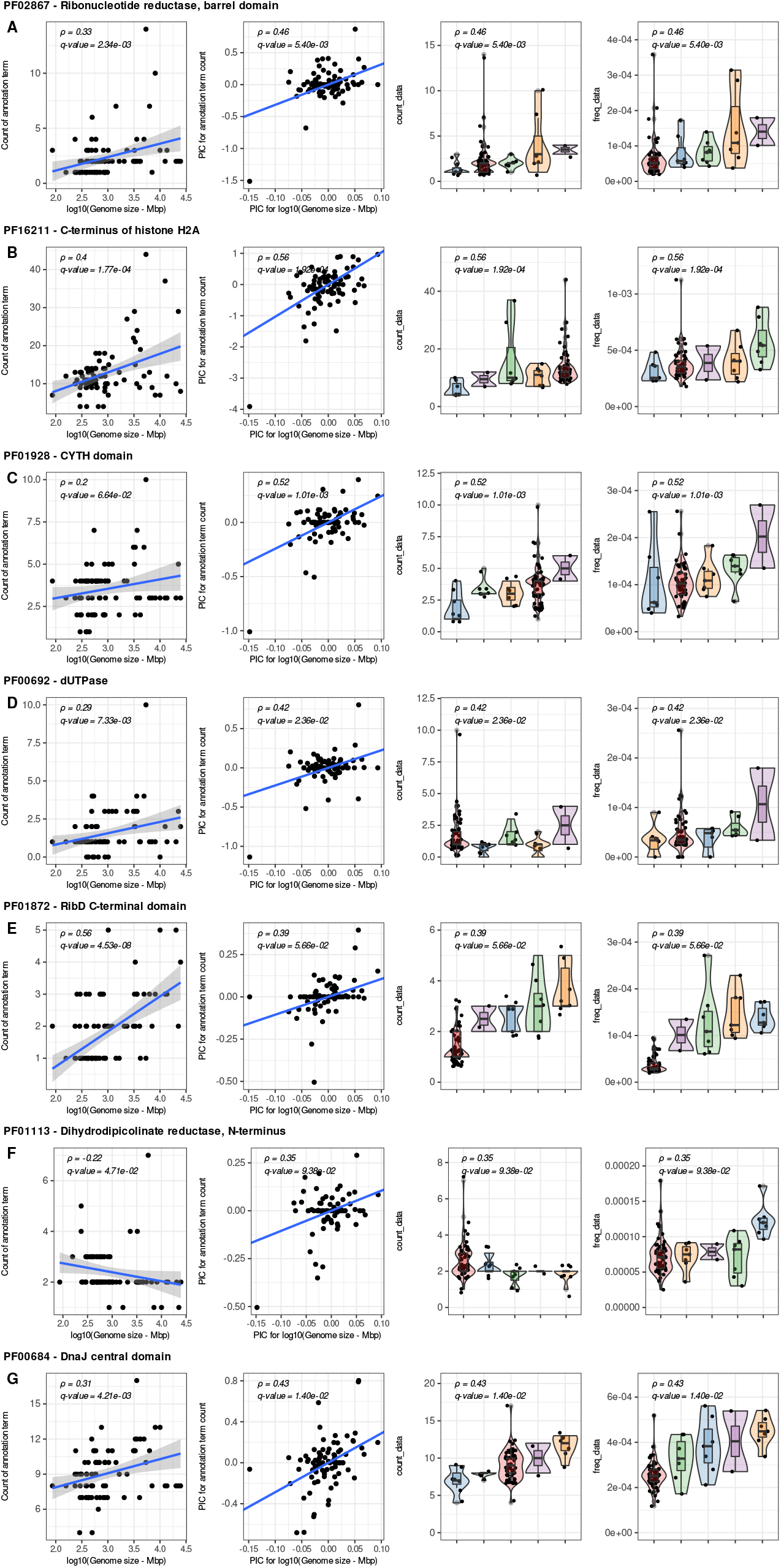
Protein domains with absolute/relative abundances in proteomes associated with genome size variation in land plants. A) Ribonucleotide reductase, barrel domain (PF02867); B) C-terminus of histone H2A (PF16211); C) CUTH domain (PF01928); D) dUTPase (PF00692); E) RibD C-terminal domain (PF01872); F) Dihydrodipicolinate reductase, N-terminus (PF01113); G) and DnaJ central domain (PF00684).

The absolute count of 3 Pfam domains was positively correlated with the increase of genome size in land plants. We also found 4 Pfam domains with relative frequencies positively associated with the increase in genome size in land plants, and a single Pfam domain (PF02867) found to be associated when considering both absolute and relative abundancies (Supplementary Table S2). Based on information mined from the manual curation of *A. thaliana* genes (Pasha et al. 2020), we noticed that as genome size increases the abundance and frequency of four Pfam domains found in proteins playing critical roles in DNA biology, such as nucleotide metabolism, DNA repair, and chromatin organization.

Two out of the six Pfam domains with positive associations are key enzymes of nucleotide metabolism: PF02867 (Ribonucleotide reductase, barrel domain) and PF00692 (dUTPase). Ribonucleotide reductases are key enzymes for DNA synthesis as they produce deoxyribonucleotides through the reduction of ribonucleotides. Deoxyuridine triphosphatases (dUTPases) are a group of enzymes that hydrolyze dUTP (deoxyuridine monophosphate) to dUMP (deoxyuridine monophosphate) and pyrophosphate. Through this reaction, dUTPases perform two key functions for DNA metabolism and repair: 1) maintain a low dUTP level in the cell, therefore preventing the misincorporation of dUTP instead of dTTP by DNA polymerase; 2) provides dUMP, a component for dTTP synthesis (Dubois et al. 2011). Therefore, the positive association of both enzymatic activities and genome size suggests these critical components of DNA metabolism and repair may play important roles for the stability of larger genomes.

The domain PF01872 (RibD C-terminal domain), also positively associated with genome size, is found in components of the riboflavin biosynthesis pathway. Riboflavin is a starting molecular scaffold for the sequential production of Flavin adenine dinucleotide (FAD) and Flavin mononucleotide (FNM). In plants, FAD and DNM are essential co-factors required for DNA repair and also for several central metabolic pathways such as photosynthesis, mitochondrial electron transport, fatty acid oxidation and photoreception (Sandoval, Y. Zhang, and Roje 2008). We hypothesize that the role of these cofactors on DNA repair may provide more genomic stability for larger genomes.

Our results indicate that as the genome size of land plants is also associated with an increase in the frequency of the C-terminus domain of the histone H2A superfamily (PF16211). The protein variants in this superfamily contain histones responsible for nucleosome formation, regulation of DNA transcription and replication, and repair of DNA double-strand breaks (Kawashima et al. 2015). In *A. thaliana*, H2A variants are also responsible for the silencing of transposable elements (TEs) (Lei and Berger 2020). It is well accepted that divergence in genome size between species is mainly due to the abundance of TEs (Bennet and Ilia J. Leitch 2005). Thus, the association of H2A protein frequency and genome size could signal the evolution of a mechanism to protect genome integrity from TEs amplification and translocation. To the best of our knowledge, these associations have not been reported elsewhere, and provide compelling evidence of how ARCADE can be used to produce new, biologically relevant knowledge.

The three remaining associations of homologous regions and genome size in land plants lack clear interpretations, and may indicate previously unknown biological processes associated with genome size variation. The CYTH domain (PF01928) converts ATP to 3’,5’-cyclic AMP and pyrophosphate. In *A. thaliana*, this domain is found in three triphosphate tunnel metalloenzymes, a poorly characterized group of homologs with roles in senescence and in defense response to pathogens (Ung, Moeder, and Yoshioka 2014). Proteins containing Pfam PF00684 (DnaJ central domain) are co-chaperones that act together with Hsp70 proteins regulating protein folding and homeostasis in response to several biotic and abiotic stresses (Liu and Whitham 2013; Pulido and Leister 2018). The only negative association was between genome size increase and PF01113 (Dihydrodipicolinate reductase, N-terminus). This domain annotates three *A. thaliana* genes involved in the diaminopimelate biosynthetic process, a crucial molecule linking the mitochondrial electron transport chain, a key component of energetic metabolism, to amino acid catabolism and the tricarboxylic acid cycle (Cavalcanti et al. 2018).

## 4 CONCLUSION

As the number of high-quality annotated genomes for major cellular lineages increases due to both the reduction of sequencing costs and the improvements of DNA sequencing technologies, genome assembly and annotation algorithms, the challenge to extract biologically meaningful, statistically sound knowledge changes from data production to data curation and modelling (Nagy et al. 2020). ARCADE was developed to address both issues by specifically providing two major data types needed for comparative genomics: sets of genomic elements shared across species’ genomes of interest and annotated to common dictionaries of biological terms and a species tree to generate phylogeny-aware models. As we demonstrated, ARCADE is the first database to provide such data types for a rich set of Archaeplastida groups not available anywhere else.

Our case study of the evolution of genome size in land plants found that species with large genome sizes have in common the presence of a higher density of genes dedicated to DNA biology. These functions may be key to preserving the stability of larger genomes. Protein variants of the histone superfamily 2A, which we found correlated with genome size increase, mark heterochromatic regions of the genome for DNA methylation silencing of TEs (Yelagandula et al. 2014). These proteins are not likely to be the reason why some plant genomes are larger, but might be a mechanism selected in lineages with larger genomes through which they cope with genome size changes. Further studies should focus on better understanding how these proteins relate to the evolution and function of plant genomes, including TEs.

## 5 DATA AVAILABILITY

All processed data needed to fully reproduce the case study (genome annotation files, phylogenetic tree, phenotypic information and CALANGO configuration files) are available at https://bit.ly/ARCADE_OSF. All raw data used in case studies (genome IDs and sources for phenotypic data) are available as supplementary tables.

## 6 ACKNOWLEDGEMENT

We would like to thank Dr. Anderson Vieira Chaves for the insightful discussions during the elaboration of this research, and Dr. Ilia J Leitch for sharing the genome size data.

## 7 FUNDING

This research had financial support from the Graduate Program in Genetics, the Graduate Program in Bioinformatics, and the Vice Dean for Research from Universidade Federal de Minas Gerais, Brazil, an d computational support from the “Centro de Processamento de Alto Desempenho/ICB” (CEPAD/SAGARANA cluster). This work was partially funded by CAPES/Brazil (Grant 001).

## 8 CONFLICT OF INTEREST

All authors declare no conflict of interest for this publication

## SUPPLEMENTARY FIGURE LEGENDS

**Figure S1: Flowchart illustrating a summarized view of the procedures performed in this project**.

**Figure S2: Distribution of sizes of non-redundant proteomes in the major Archaeplastida clades after each one of the methods we use for redundancy reductions (CD-HIT (W. Li and Godzik 2006; Fu et al. 2012) and our in-house pipeline)**.

**Figure S3: Comparison of CD-HIT (W. Li and Godzik 2006; Fu et al. 2012) and our in-house method for redundancy reduction according to BUSCO’s (Manni et al. 2021) gene completeness (C)**.

**Figure S4: Comparison of CD-HIT (W. Li and Godzik 2006; Fu et al. 2012) and our in-house method for redundancy reduction according to BUSCO’s (Manni et al. 2021) single-copy genes (S)**.

**Figure S5: Comparison of CD-HIT (W. Li and Godzik 2006; Fu et al. 2012) and our in-house method for redundancy reduction according to BUSCO’s (Manni et al. 2021) gene ducplication (D)**.

**Figure S6: Comparison of CD-HIT (W. Li and Godzik 2006; Fu et al. 2012) and our in-house method for redundancy reduction according to BUSCO’s (Manni et al. 2021) fragmented genes (F)**.

**Figure S7: Comparison of CD-HIT (W. Li and Godzik 2006; Fu et al. 2012) and our in-house method for redundancy reduction according to BUSCO’s (Manni et al. 2021) missing gene (M)**.

**Table S1:** List of species we surveyed and their respective clade, species code, and log10 transformed genome size. The full table with information on the database of origin, clade, BUSCO (Manni et al. 2021) results, genome size in picograms, megabase pairs, log10 transformed, and genome ID is available at https://bit.ly/ARCADE_tableS1_FULL.

| Clade | Species | Code | log10(GS) |
| --- | --- | --- | --- |
| Angiosperm | *Acer yangbiense* | Aya | NA |
| Angiosperm | *Akebia trifoliata* | Atr | 2.8711 |
| Angiosperm | *Amaranthus hypochondriacus* | Ahy | 2.6893 |
| Angiosperm | *Amborella trichopoda* | Ati | 2.9393 |
| Angiosperm | *Ananas comosus* | Aco | 2.7276 |
| Angiosperm | *Arabidopsis halleri* | Aha | 2.3705 |
| Angiosperm | *Arabidopsis thaliana* | Ath | 2.1953 |
| Angiosperm | *Arabis nemorensis* | Ane | NA |
| Angiosperm | *Ascarina rubricaulis* | Arb | NA |
| Angiosperm | *Asparagus officinalis* | Aof | 3.1555 |
| Angiosperm | *Beta vulgaris subsp. vulgaris* | Bvu | 2.955 |
| Angiosperm | *Boechera stricta* | Bst | 2.3705 |
| Angiosperm | *Brachypodium distachyon* | Bdi | 2.4902 |
| Angiosperm | *Brachypodium stacei* | Bsa | NA |
| Angiosperm | *Brachypodium sylvaticum* | Bsy | 2.6238 |
| Angiosperm | *Cajanus cajan* | Cca | 2.9454 |
| Angiosperm | *Cannabis sativa* | Csa | 2.9146 |
| Angiosperm | *Capsicum annuum* | Can | 3.4909 |
| Angiosperm | *Capsicum baccatum* | Cba | 3.5605 |
| Angiosperm | *Capsicum chinense* | Cch | 3.5252 |
| Angiosperm | *Carex littledalei* | Cli | NA |
| Angiosperm | *Carica papaya* | Cpa | 2.5932 |
| Angiosperm | *Castanea mollissima* | Cmo | NA |
| Angiosperm | *Cephalotus follicularis* | Cfo | 2.7957 |
| Angiosperm | *Cinnamomum kanehirae* | Cka | NA |
| Angiosperm | *Cinnamomum micranthum* | Cmi | NA |

Table S1 continued from previous page
| Clade | Species | Code | log10(GS) |
| --- | --- | --- | --- |
| Angiosperm | *Citrus clementina* | Ccl | NA |
| Angiosperm | *Citrus sinensis* | Csi | 2.7693 |
| Angiosperm | *Citrus unshiu* | Cun | NA |
| Angiosperm | *Cleome violacea* | Cvi | NA |
| Angiosperm | *Corchorus capsularis* | Ccp | NA |
| Angiosperm | *Corchorus olitorius* | Col | NA |
| Angiosperm | *Corymbia citriodora* | Cci | 2.5814 |
| Angiosperm | *Cucumis melo* | Cme | 2.9912 |
| Angiosperm | *Cucumis melo* var. *makuwa* | Cmm | NA |
| Angiosperm | *Cucumis sativus* | Cst | 2.9454 |
| Angiosperm | *Cuscuta australis* | Cau | NA |
| Angiosperm | *Dendrobium officinale* | Dof | 3.6236 |
| Angiosperm | *Descurainia sophioides* | Dso | NA |
| Angiosperm | *Dioscorea alata* | Dal | 2.7537 |
| Angiosperm | *Doroceras hygrometricum* | Dhy | NA |
| Angiosperm | *Erythranthe guttata* | Egt | 2.5585 |
| Angiosperm | *Eucalyptus grandis* | Egr | 2.7871 |
| Angiosperm | *Fragaria vesca* subsp. *vesca* | Fve | 2.3821 |
| Angiosperm | *Hakea prostrata* | Hpo | NA |
| Angiosperm | *Herrania umbratica* | Hum | NA |
| Angiosperm | *Hibbertia grossulariifolia* | Hgr | NA |
| Angiosperm | *Hordeum vulgare* | Hvu | 3.7307 |
| Angiosperm | *Hypocoum procumbens* | Hpr | NA |
| Angiosperm | *Illicium floridanum* | Ifl | NA |
| Angiosperm | *Jatropha curcas* | Jcu | 2.6196 |
| Angiosperm | *Kingdonia uniflora* | Kun | NA |
| Angiosperm | *Lactuca saligna* | Lsa | 3.3669 |
| Angiosperm | *Lactuca sativa* | Lst | 3.5103 |
| Angiosperm | *Macleaya cordata* | Mco | NA |
| Angiosperm | *Momordica charantia* | Mch | 3.3134 |

Table S1 continued from previous page
| Clade | Species | Code | log10(GS) |
| --- | --- | --- | --- |
| Angiosperm | *Morus notabilis* | Mno | NA |
| Angiosperm | *Musa balbisiana* | Mba | 2.7684 |
| Angiosperm | *Nymphaea colorata* | Nco | 2.6893 |
| Angiosperm | *Nymphaea thermarum* | Nth | 2.6979 |
| Angiosperm | *Oropetium thomaeum* | Oth | NA |
| Angiosperm | *Oryza brachyantha* | Obr | 2.5932 |
| Angiosperm | *Oryza meyeriana* var. *granulata* | Ome | 2.9445 |
| Angiosperm | *Oryza sativa* | Osa | NA |
| Angiosperm | *Oryza sativa* Indica Group | Osi | NA |
| Angiosperm | *Oryza sativa* Japonica Group | Osj | 2.6901 |
| Angiosperm | *Parasponia andersonii* | Pan | NA |
| Angiosperm | *Phalaenopsis equestris* | Peq | 3.2178 |
| Angiosperm | *Phaseolus lunatus* | Plu | 2.8354 |
| Angiosperm | *Phaseolus vulgaris* | Pvu | 2.7693 |
| Angiosperm | *Podophyllum peltatum* | Ppe | NA |
| Angiosperm | *Prunus armeniaca* | Par | NA |
| Angiosperm | *Prunus avium* | Pav | 2.6246 |
| Angiosperm | *Prunus dulcis* | Pdu | 2.5088 |
| Angiosperm | *Prunus mume* | Pmu | NA |
| Angiosperm | *Prunus persica* | Ppr | 2.4683 |
| Angiosperm | *Punica granatum* | Pgr | 2.8485 |
| Angiosperm | *Rhamnella rubrinervis* | Rru | NA |
| Angiosperm | *Rhododendron williamsianum* | Rwi | NA |
| Angiosperm | *Ricinus communis* | Rco | 2.7075 |
| Angiosperm | *Solanum lycopersicum* | Sly | 3.0165 |
| Angiosperm | *Solanum pennellii* | Spe | 3.0837 |
| Angiosperm | *Solanum tuberosum* | Stu | 2.9381 |
| Angiosperm | *Sorghum bicolor* | Sbi | 2.8363 |
| Angiosperm | *Spatholobus suberectus* | Ssu | NA |
| Angiosperm | *Spinacia oleracea* | Sol | 2.9912 |

Table S1 continued from previous page
| Clade | Species | Code | log10(GS) |
| --- | --- | --- | --- |
| Angiosperm | *Spirodela polyrhiza* | Spo | 2.4674 |
| Angiosperm | *Syzygium oleosum* | Soe | NA |
| Angiosperm | *Thalictrum thalictroides* | Tth | 3.5538 |
| Angiosperm | *Theobroma cacao* | Tca | 2.5932 |
| Angiosperm | *Trema orientale* | Tor | NA |
| Angiosperm | *Trifolium subterraneum* | Tsu | 2.7355 |
| Angiosperm | *Vigna angularis* | Van | 2.6901 |
| Angiosperm | *Vigna radiata* var. *radiata* | Vra | 2.6901 |
| Angiosperm | *Vigna unguiculata* | Vun | 2.7684 |
| Angiosperm | *Vitis riparia* | Vri | 2.6893 |
| Angiosperm | *Vitis vinifera* | Vvi | 2.5932 |
| Angiosperm | *Zostera marina* | Zmr | 2.4895 |
| Bryophyta | *Andreaea rupestris* | Aru | 2.3125 |
| Bryophyta | *Anthoceros angustus* | Aan | NA |
| Bryophyta | *Aulacomnium heterostichum* | Ahe | 2.6435 |
| Bryophyta | *Buxbaumia aphylla* | Bap | NA |
| Bryophyta | *Ceratodon purpureus* | Cpu | 2.5814 |
| Bryophyta | *Claopodium rostratum* | Cro | NA |
| Bryophyta | *Diphyscium foliosum* | Dfo | NA |
| Bryophyta | *Frisvollia varia* | Fva | NA |
| Bryophyta | *Frullania* spp. | Fru | NA |
| Bryophyta | *Hedwigia ciliata* | Hci | 2.4674 |
| Bryophyta | *Leucobryum albidum* | Lal | NA |
| Bryophyta | *Lunularia cruciata* | Lcr | 2.8164 |
| Bryophyta | *Marchantia polymorpha* | Mpo | 2.458 |
| Bryophyta | *Marchantia polymorpha* subsp. *ruderalis* | Mpr | NA |
| Bryophyta | *Neckera douglasii* | Ndo | NA |
| Bryophyta | *Pellia neesiana* | Pne | NA |
| Bryophyta | *Philonotis fontana* | Pfo | NA |
| Bryophyta | *Porella navicularis* | Pna | 2.9146 |

Table S1 continued from previous page
| Clade | Species | Code | log10(GS) |
| --- | --- | --- | --- |
| Bryophyta | *Pseudanomodon attenuatus* | Pat | 2.5923 |
| Bryophyta | *Pseudotaxiphyllum elegans* | Pel | NA |
| Bryophyta | *Pulviger a lyellii* | Ply | NA |
| Bryophyta | *Ricciocarpos natans* | Rna | NA |
| Bryophyta | *Takakia lepidozoides* | Tle | NA |
| Bryophyta | *Timmia austriaca* | Tau | NA |
| Charophyta | *Chara braunii* | Cbr | NA |
| Charophyta | *Coleochaete irregularis* | Cir | NA |
| Charophyta | *Cylindrocystis cushleackae* | Ccu | NA |
| Charophyta | *Klebsormidium nitens* | Kni | NA |
| Charophyta | *Mesotaenium caldariorum* | Mca | NA |
| Charophyta | *Staurodesmus omearii* | Som | NA |
| Chlorophyta | *Auxenochlorella protothecoides* | Apr | NA |
| Chlorophyta | *Chlamydomonas eustigma* | Ceu | NA |
| Chlorophyta | *Chlamydomonas reinhardtii* | Cre | NA |
| Chlorophyta | *Chlorella sorokiniana* | Cso | NA |
| Chlorophyta | *Chlorella variabilis* | Cva | NA |
| Chlorophyta | *Chloropicon primus* | Cpr | NA |
| Chlorophyta | *Chromochloris zofingiensis* | Czo | NA |
| Chlorophyta | *Coccomyxa subellipsoidea C-169* | Csu | NA |
| Chlorophyta | *Micractinium conductrix* | Mcn | NA |
| Chlorophyta | *Micromonas commoda* | Mcm | NA |
| Chlorophyta | *Micromonas pusilla CCMP1545* | Mpu | NA |
| Chlorophyta | *Ostreococcus tauri* | Ota | NA |
| Chlorophyta | *Raphidocelis subcapitata* | Rsu | NA |
| Chlorophyta | *Scenedesmus sp. NREL 46B-D3* | Sce | NA |
| Chlorophyta | *Trebouxia sp. A1-2* | Tre | NA |
| Chlorophyta | *Volvox carteri f. nagariensis* | Vca | NA |
| Glaucophyta | *Gloeochaete wittrockiana* | Gwi | NA |
| Gymnosperm | *Cephalotaxus harringtonia* | Cha | 4.3943 |

Table S1 continued from previous page
| Clade | Species | Code | log10(GS) |
| --- | --- | --- | --- |
| Gymnosperm | *Gnetum montanum* | Gmo | NA |
| Gymnosperm | *Microcachrys tetragona* | Mte | 3.6083 |
| Gymnosperm | *Pinus sylvestris* | Psy | 4.3525 |
| Gymnosperm | *Sciadopitys verticillata* | Sve | 4.3084 |
| Gymnosperm | *Thuja plicata* | Tpl | 4.0989 |
| Gymnosperm | *Welwitschia mirabilis* | Wmi | 3.8476 |
| Gymnosperm | *Wollemia nobilis* | Wno | 4.1346 |
| Lycophyta | *Huperzia selago* | Hse | 3.6936 |
| Lycophyta | *Isoetes tegetiformans* | Ite | NA |
| Lycophyta | *Selaginella moellendorffii* | Smo | 1.9357 |
| Monilophyta | *Asplenium platyneuron* | Apl | NA |
| Monilophyta | *Azolla filiculoides* | Afi | 2.8767 |
| Monilophyta | *Cystopteris fragilis* | Cfr | 3.9088 |
| Monilophyta | *Danaea nodosa* | Dno | NA |
| Monilophyta | *Diplazium wichurae* | Dwi | NA |
| Monilophyta | *Leucostegia immersa* | Lim | NA |
| Monilophyta | *Lonchitis hirsuta* | Lhi | NA |
| Monilophyta | *Pilularia globulifera* | Pgl | 3.0073 |
| Monilophyta | *Plenasium javanicum* | Pja | NA |
| Monilophyta | *Polypodium glycyrrhiza* | Pgy | 4.001 |
| Monilophyta | *Salvinia cucullata* | Scu | 2.4065 |
| Monilophyta | *Sceptridium dissectum* | Sdi | 3.5256 |
| Monilophyta | *Struthiopteris spicant* | Ssp | 3.7992 |
| Monilophyta | *Vittaria appalachiana* | Vap | NA |
| Prasinodermophyta | *Prasinoderma coloniale* | Pcl | NA |
| Rhodophyta | *Galdieria sulphuraria* | Gsu | NA |

**Table S2:** List of *Arabidopsis thaliana* genes containing the proteins domains found correlated to genome size evolution in land plants by our analysis with their standard identifications and annotation.

| Corr | Pfam | gene symbol | locus tag | gene name | Short description from TAIR |
| --- | --- | --- | --- | --- | --- |
| negative | PFO1113 | AT2G44040 | AT2G44040 | - | Dihydrodipicolinate reductase, bacterial/plant |
| negative | PFO1113 | AT3G59890 | AT3G59890 | - | Dihydrodipicolinate reductase, bacterial/plant |
| negative | PFO1113 | AT5G52100 | AT5G52100 | CRR1 | Dihydrodipicolinate reductase, bacterial/plant |
| positive | PF00684 | AT1G28210 | AT1G28210 | ATJ1 | DNAJ heat shock family protein |
| positive | PF00684 | AT1G80030 | AT1G80030 | DJA7 | Molecular chaperone Hsp40/DnaJ family protein |
| positive | PF00684 | AT2G22360 | AT2G22360 | DJA6 | DNAJ heat shock family protein |
| positive | PF00684 | AT3G17830 | AT3G17830 | DJA4 | Molecular chaperone Hsp40/DnaJ family protein |
| positive | PF00684 | AT3G44110 | AT3G44110 | J3 | DNAJ homologue 3 |
| positive | PF00684 | AT4G39960 | AT4G39960 | DJA5 | Molecular chaperone Hsp40/DnaJ family protein |
| positive | PF00684 | J2 | AT5G22060 | J2 | DNAJ homologue 2 |
| positive | PF00684 | GFA2 | AT5G48030 | GFA2 | gametophytic factor 2 |
| positive | PF00692 | AT3G46940 | AT3G46940 | DUT1 | DUTP-PYROPHOSPHATASE-LIKE 1 |
| positive | PF01872 | AT3G47390 | AT3G47390 | PHS1 | cytidine/deoxycytidylate deaminase family protein |
| positive | PF01928 | AT1G26190 | AT1G26190 | TTM2 | Phosphoribulokinase / Uridine kinase family |
| positive | PF01928 | AT1G73980 | AT1G73980 | TTM1 | Phosphoribulokinase / Uridine kinase family |
| positive | PF01928 | AT2G11890 | AT2G11890 | TTM3 | adenylate cyclases |
| positive | PF02867 | AT2G21790 | AT2G21790 | RNR1 | ribonucleotide reductase 1 |
| positive | PF16211 | AT1G08880 | AT1G08880 | H2AXA | histone superfamily protein |
| positive | PF16211 | AT1G51060 | AT1G51060 | HTA10 | histone H2A 10 |
| positive | PF16211 | HTA9 | AT1G52740 | HTA9 | histone H2A protein 9 |
| positive | PF16211 | AT1G54690 | AT1G54690 | HTA3 | gamma histone variant H2AX |
| positive | PF16211 | AT2G38810 | AT2G38810 | HTA8 | histone H2A 8 |
| positive | PF16211 | HTA13 | AT3G20670 | HTA13 | histone H2A 13 |
| positive | PF16211 | AT3G54560 | AT3G54560 | HTA11 | histone H2A 11 |
| positive | PF16211 | AT4G27230 | AT4G27230 | HTA2 | histone H2A 2 |
| positive | PF16211 | AT5G02560 | AT5G02560 | HTA12 | histone H2A 12 |
| positive | PF16211 | AT5G27670 | AT5G27670 | HTA7 | histone H2A 7 |
| positive | PF16211 | AT5G54640 | AT5G54640 | RAT5 | histone superfamily protein |
| positive | PF16211 | AT5G59870 | AT5G59870 | HTA6 | histone H2A 6 |

## Notes

https://osf.io/2fkvh/

### Competing Interest Statement

The authors have declared no competing interest.

## REFERENCES

Adams, Dean C. and Michael L. Collyer (July 2017). “Multivariate Phylogenetic Comparative: Evaluations, Comparisons, and Recommendations”. Systematic Biology 67.1, pp. 14–31.

Beaulieu, Jeremy M. et al. (2007). “Correlated evolution of genome size and seed mass”. New Phytologist 173.2, pp. 422–437.

Bennet, Michael D. and Ilia J. Leitch (2005). “CHAPTER 2 - Genome Size Evolution in Plants”. The Evolution of the Genome. Ed. by T. Ryan Gregory. Burlington: Academic Press, pp. 89–162. isBn: 978-0-12-301463-4

Bennett, Michael D. (1987). “VARIATION IN GENOMIC FORM IN PLANTS AND ITS ECOLOGICAL IMPLICATIONS”. New Phytologist 106.1, pp. 177–200.

Blázquez, Miguel A., David C. Nelson, and Dolf Weijers (2020). “Evolution of Plant Hormone Response Pathways”. Annual Review of Plant Biology 71.1. pp. 327–353.

Carta, Angelino et al. (2022). “Correlated evolution of seed mass and genome size varies among life forms in flowering plants”. Seed Science Research 32.1, pp. 46–52.

Cavalcanti, João Henrique F et al. (Sept. 2018). “An L,L-diaminopimelate aminotransferase mutation leads to metabolic shifts and growth inhibition in Arabidopsis”. Journal of Experimental Botany 69.22, pp. 5489–5506.

Challis, Richard J. et al. (May 2017). “GenomeHubs: simple containerized setup of a custom Ensembl database and web server for any species”. Database 2017. bax039.

Chan, Patricia P. and Todd M. Lowe (Dec. 2015). “GtRNAdb 2.0: an expanded database of transfer RNA genes identified in complete and draft genomes”. Nucleic Acids Research 44.D1, pp. D184–D189. issn: 0305-1048.

Chanderbali, Andre S et al. (Mar. 2016). “Evolving Ideas on the Origin and Evolution of Flowers: New Perspectives in the Genomic Era”. Genetics 202.4, pp. 1255–1265.

Chen, Lu et al. (Mar. 2014). “Correcting for Differential Transcript Coverage Reveals a Strong Relationship between Alternative Splicing and Organism Complexity”. Molecular Biology and Evolution 31.6, pp. 1402–1413.

Dubois, Emeline et al. (Apr. 2011). “Homologous Recombination Is Stimulated by a Decrease in dUTPase in Arabidopsis”. PLOS ONE 6.4, pp. 1–8.

Ekstrom, Alexander et al. (Aug. 2014). “PlantCAZyme: a database for plant carbohydrate-active enzymes”. Database 2014. bau079. issn: 1758-0463.

Felsenstein, Joseph (1985). “Phylogenies and the Comparative Method”. The American Naturalist 125.1, pp. 1–15.

Fleischmann, Andreas et al. (Oct. 2014). “Evolution of genome size and chromosome number in the carnivorous plant genus Genlisea (Lentibulariaceae), with a new estimate of the minimum genome size in angiosperms”. Annals of Botany 114.8, pp. 1651– 1663.

Fu, Limin et al. (Oct. 2012). “CD-HIT: accelerated for clustering the next-generation sequencing data”. Bioinformatics 28.23, pp. 3150–3152.

Goodstein, David M. et al. (Nov. 2011). “Phytozome: a comparative platform for green plant genomics”. Nucleic Acids Research 40.D1, pp. D1178–D1186.

Hidalgo, Oriane et al. (2017). “Is There an Upper Limit to Genome Size?” Trends in Plant Science 22.7, pp. 567–573.

Hongo, Jorge Augusto et al. (2021). “CALANGO: an annotation-based, phylogeny-aware comparative genomics framework for exploring and interpreting complex genotypes and phenotypes”. bioRxiv. doi: 10.1101/2021.08.25.457574.

Jones, Philip et al. (Jan. 2014). “InterProScan 5: genome-scale protein function classification”. Bioinformatics 30.9, pp. 1236–1240.

Kang, Ming et al. (2014). “Adaptive and nonadaptive genome size evolution in Karst endemic flora of China”. New Phytologist 202.4, pp. 1371–1381.

Kattge, Jens et al. (2020). “TRY plant trait database – enhanced coverage and open access”. Global Change Biology 26.1, pp. 119–188.

Kawashima, Tomokazu et al. (July 2015). “Diversification of histone H2A variants during plant evolution”. Trends in Plant Science 20.7, pp. 419–425.

Knight, Charles A., Nicole A. Molinari, and Dmitri A. Petrov (Jan. 2005). “The Large Genome Constraint Hypothesis: Evolution, Ecology and Phenotype”. Annals of Botany 95.1, pp. 177–190.

Krishnakumar, Vivek et al. (Nov. 2014). “Araport: the Arabidopsis Information Portal”. Nucleic Acids Research 43.D1, pp. D1003–D1009.

Kumar, Sudhir et al. (Aug. 2022). “TimeTree 5: An Expanded Resource for Species Divergence Times”. Molecular Biology and Evolution 39.8. msac174.

Lei, Bingkun and Frédéric Berger (2020). “H2A Variants in Arabidopsis: Versatile Regulators of Genome Activity”. Plant Communications 1.1, p. 100015.

Leitch, A. R. and I. J. Leitch (2012). “Ecological and genetic factors linked to contrasting genome dynamics in seed plants”. New Phytologist 194.3, pp. 629–646.

Leitch, I. J. et al. (Jan. 2005). “Evolution of DNA Amounts Across Land Plants (Embryophyta)”. Annals of Botany 95.1, pp. 207–217.

Leitch, Ilia J., Mark W. Chase, and Michael D. Bennett (Dec. 1998). “Phylogenetic Analysis of DNA C-values Provides Evidence for a Small Ancestral Genome Size in Flowering Plants”. Annals of Botany 82.Suppl_1, pp. 85–94.

Li, Fay-Wei et al. (July 2018). “Fern genomes elucidate land plant evolution and cyanobacterial symbioses”. Nature Plants 4.7, pp. 460–472.

Li, Weizhong and Adam Godzik (May 2006). “Cd-hit: a fast program for clustering and comparing large sets of protein or nucleotide sequences”. Bioinformatics 22.13, pp. 1658–1659.

Liu, Jian-Zhong and Steven A. Whitham (2013). “Overexpression of a soybean nuclear localized type–III DnaJ domain-containing HSP40 reveals its roles in cell death and disease resistance”. The Plant Journal 74.1, pp. 110–121.

Ma, Xuelian et al. (Sept. 2022). “PlantGSAD: a comprehensive gene set annotation database for plant species”. Nucleic Acids Research 50.D1, pp. D1456–D1467. issn: 0305-1048.

Manni, Mosè et al. (July 2021). “BUSCO Update: Novel and Streamlined Workflows along with Broader and Deeper Phylogenetic Coverage for Scoring of Eukaryotic, Prokaryotic, and Viral Genomes”. Molecular Biology and Evolution 38.10, pp. 4647–4654.

Marks, Rose A. et al. (Dec. 2021). “Representation and participation across 20 years of plant genome sequencing”. Nature Plants 7.12, pp. 1571–1578.

Nagy, László G et al. (Jan. 2020). “Novel phylogenetic methods are needed for understanding gene function in the era of mega-scale genome sequencing”. Nucleic Acids Research 48.5, pp. 2209–2219.

O’Leary, Nuala A. et al. (Nov. 2015). “Reference sequence (RefSeq) database at NCBI: current status, taxonomic expansion, and functional annotation”. Nucleic Acids Research 44.D1, pp. D733–D745.

Pasha, Asher et al. (July 2020). “Araport Lives: An Updated Framework for Arabidopsis Bioinformatics”. The Plant Cell 32.9, pp. 2683–2686.

Pellicer, Jaume, Michae F. Fay, and Ilia J. Leitch (Sept. 2010). “The largest eukaryotic genome of them all?” Botanical Journal of the Linnean Society 164.1, pp. 10–15.

Pellicer, Jaume, Oriane Hidalgo, et al. (2018). “Genome Size Diversity and Its Impact on the Evolution of Land Plants”. Genes 9.2.

Pellicer, Jaume and Ilia J. Leitch (2020). “The Plant DNA C-values database (release 7.1): an updated online repository of plant genome size data for comparative studies”. New Phytologist 226.2, pp. 301–305.

Petrov, Dmitri A. (Jan. 2001). “Evolution of genome size: new approaches to an old problem”. Trends in Genetics 17.1, pp. 23–28.

Petrov, Dmitri A (2002). “Mutational Equilibrium Model of Genome Size Evolution”. Theoretical Population Biology 61.4, pp. 531–544.

Proost, Sebastian et al. (Oct. 2014). “PLAZA 3.0: an access point for plant comparative genomics”. Nucleic Acids Research 43.D1, pp. D974–D981.

Pulido, Pablo and Dario Leister (2018). “Novel DNAJ-related proteins in Arabidopsis thaliana”. The New Phytologist 217.2, pp. 480–490.

Puttick, Mark N., James Clark, and Philip C. J. Donoghue (2015). “Size is not everything: rates of genome size evolution, not C-value, correlate with speciation in angiosperms”. Proceedings of the Royal Society B: Biological Sciences 282.1820, p. 20152289.

Revell, Liam J. (2012). “phytools: an R package for phylogenetic comparative biology (and other things)”. Methods in Ecology and Evolution 3.2, pp. 217–223.

Sandoval, Francisco J., Yi Zhang, and Sanja Roje (Nov. 2008). “Flavin Nucleotide Metabolism in Plants: MONOFUNCTIONAL ENZYMES SYNTHESIZE FAD IN PLASTIDS *”. Journal of Biological Chemistry 283.45, pp. 30890–30900.

Sayers, Eric W et al. (Oct. 2019). “GenBank”. Nucleic Acids Research 48.D1, pp. D84– D86.

Shapiro, S. S. and M. B. Wilk (Dec. 1965). “An analysis of variance test for normality (complete samples)”. Biometrika 52.3-4, pp. 591–611.

Sievert, Carson (2016). ggplot2: Elegant Graphics for Data Analysis. Springer-Verlag New York. isBn: 978-3-319-24277-4.

Tello-Ruiz, Marcela K et al. (Nov. 2020). “Gramene 2021: harnessing the power of comparative genomics and pathways for plant research”. Nucleic Acids Research 49.D1, pp. D1452–D1463.

Ung, Huoi, Wolfgang Moeder, and Keiko Yoshioka (Sept. 2014). “Arabidopsis Triphosphate Tunnel Metalloenzyme2 Is a Negative Regulator of the Salicylic Acid-Mediated Feedback Amplification Loop for Defense Responses”. Plant Physiology 166.2, pp. 1009–1021.

Veselý, Pavel, Petr Bureš, and Petr Šmarda (Aug. 2013). “Nutrient reserves may allow for genome size increase: evidence from comparison of geophytes and their sister non-geophytic relatives”. Annals of Botany 112.6, pp. 1193–1200.

Vinogradov, Alexander E (2003). “Selfish DNA is maladaptive: evidence from the plant Red List”. Trends in Genetics 19.11, pp. 609–614.

Vogel, Christine and Cyrus Chothia (May 2006). “Protein Family Expansions and Biological Complexity”. PLOS Computational Biology 2.5, pp. 1–13.

Wang, Dandan et al. (2021). “Which factors contribute most to genome size variation within angiosperms?” Ecology and Evolution 11.6, pp. 2660–2668.

Wang, Xiaoxue et al. (Dec. 2010). “Cryptic prophages help bacteria cope with adverse environments”. Nature Communications 1.1, p. 147.

Wendel, Jonathan F. et al. (May 2002). “Feast and famine in plant genomes”. Genetica 115.1, pp. 37–47.

Yang, Xiaohan et al. (Sept. 2019). “Comparative genomics can provide new insights into the evolutionary mechanisms and gene function in CAM plants”. Journal of Experimental Botany 70.22, pp. 6539–6547.

Yelagandula, Ramesh et al. (July 2014). “The Histone Variant H2A.W Defines Heterochromatin and Promotes Chromatin Condensation in Arabidopsis”. Cell 158.1, pp. 98–109.

Zhang, Jian et al. (Feb. 2020). “The hornwort genome and early land plant evolution”. Nature Plants 6.2, pp. 107–118.

