## Supplementary figures and images for "Using ARCADE (ARChaeplastida Annotation DatabasE) to understand the evolution of genome size in land plants"

### Fig. S1

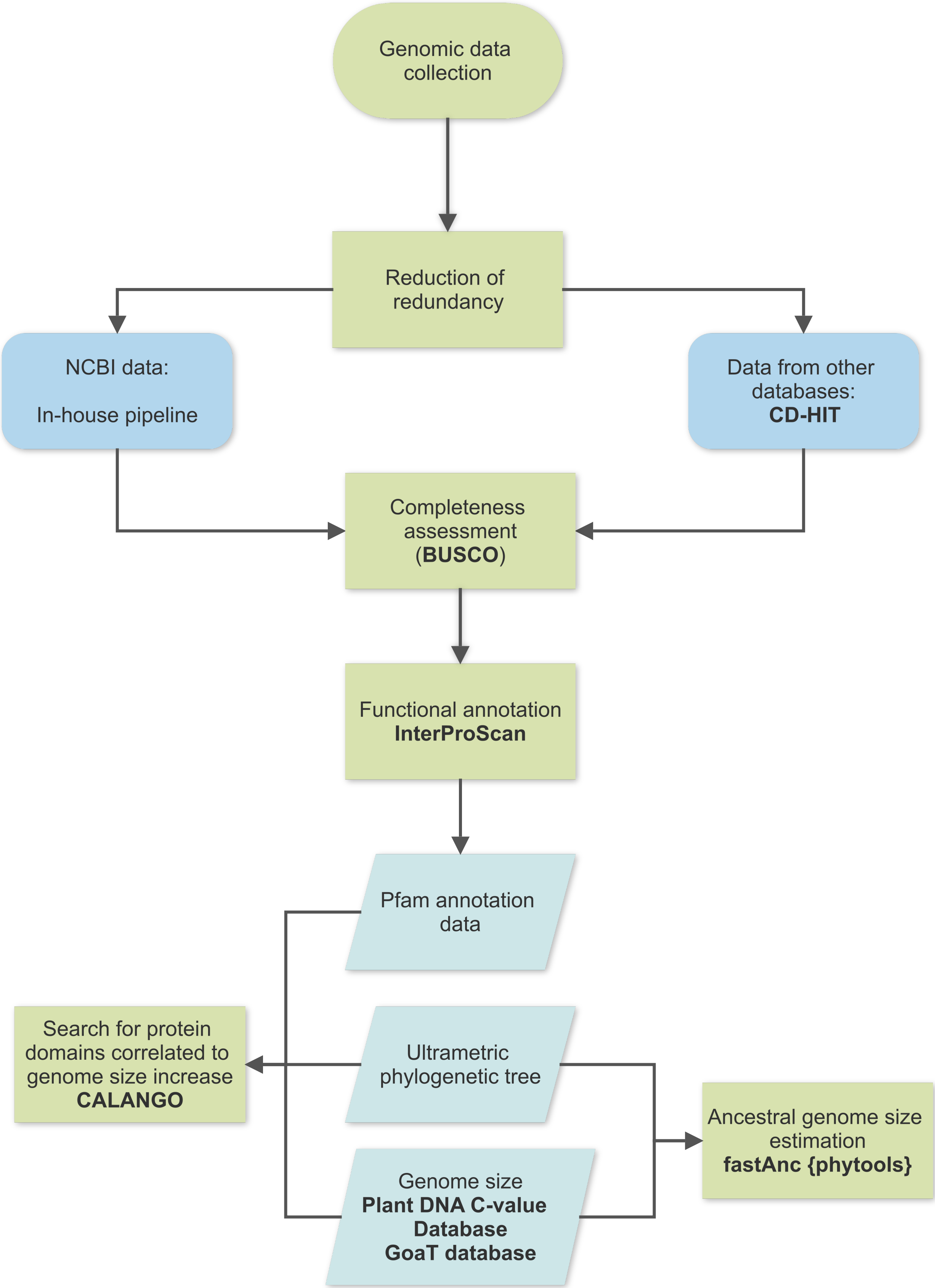

### Fig. S2

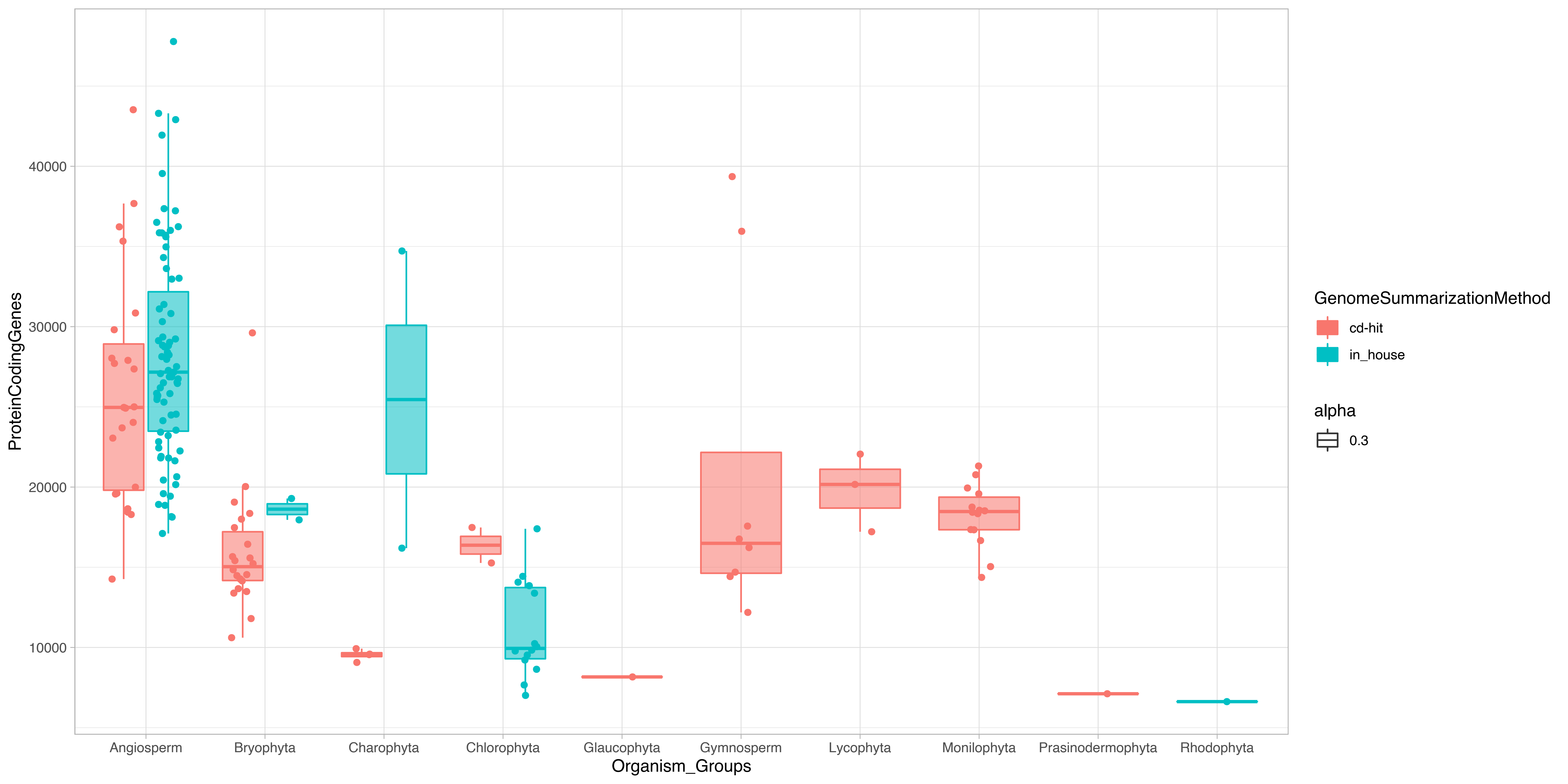

### Fig. S3

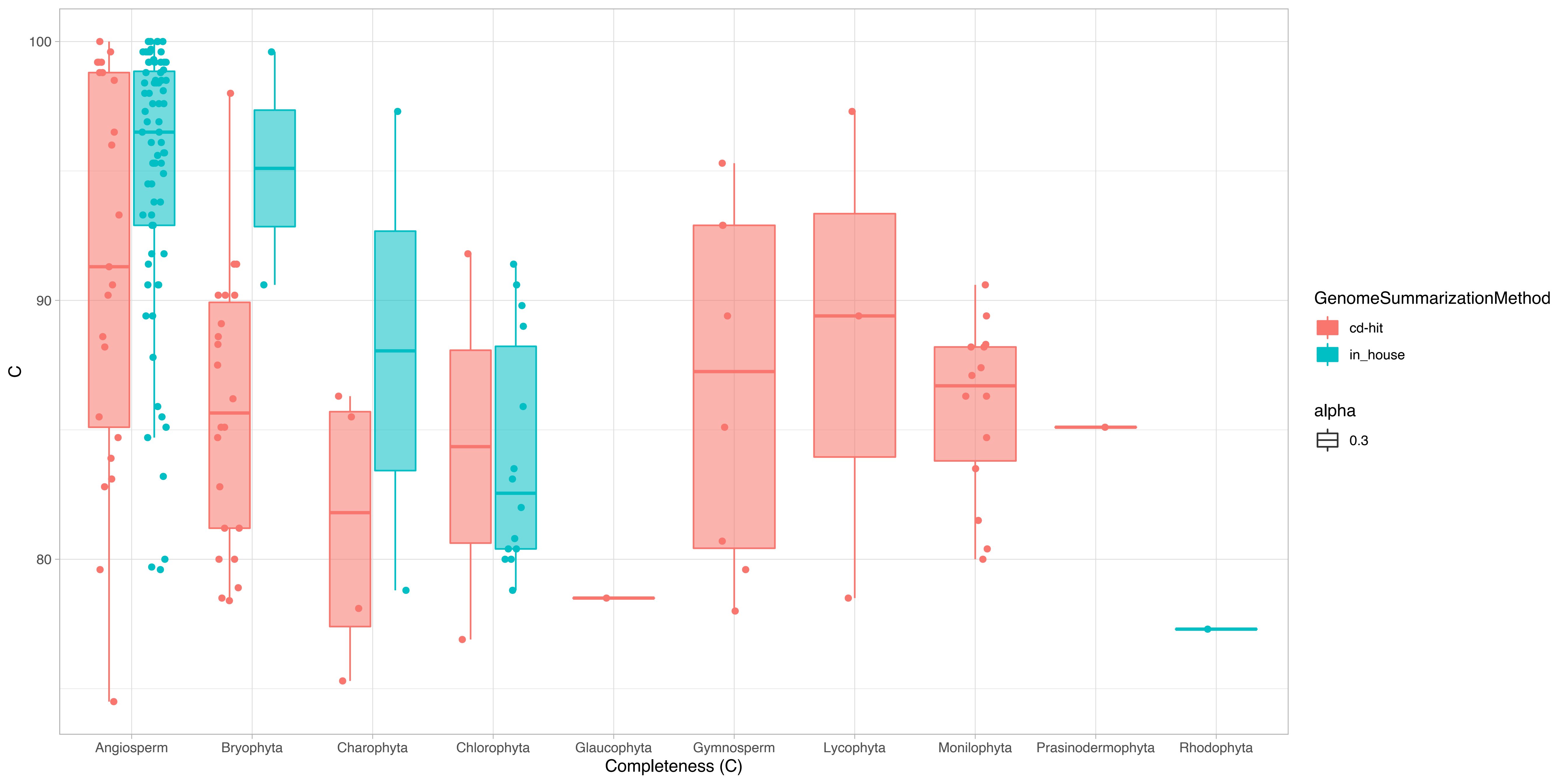

### Fig. S4

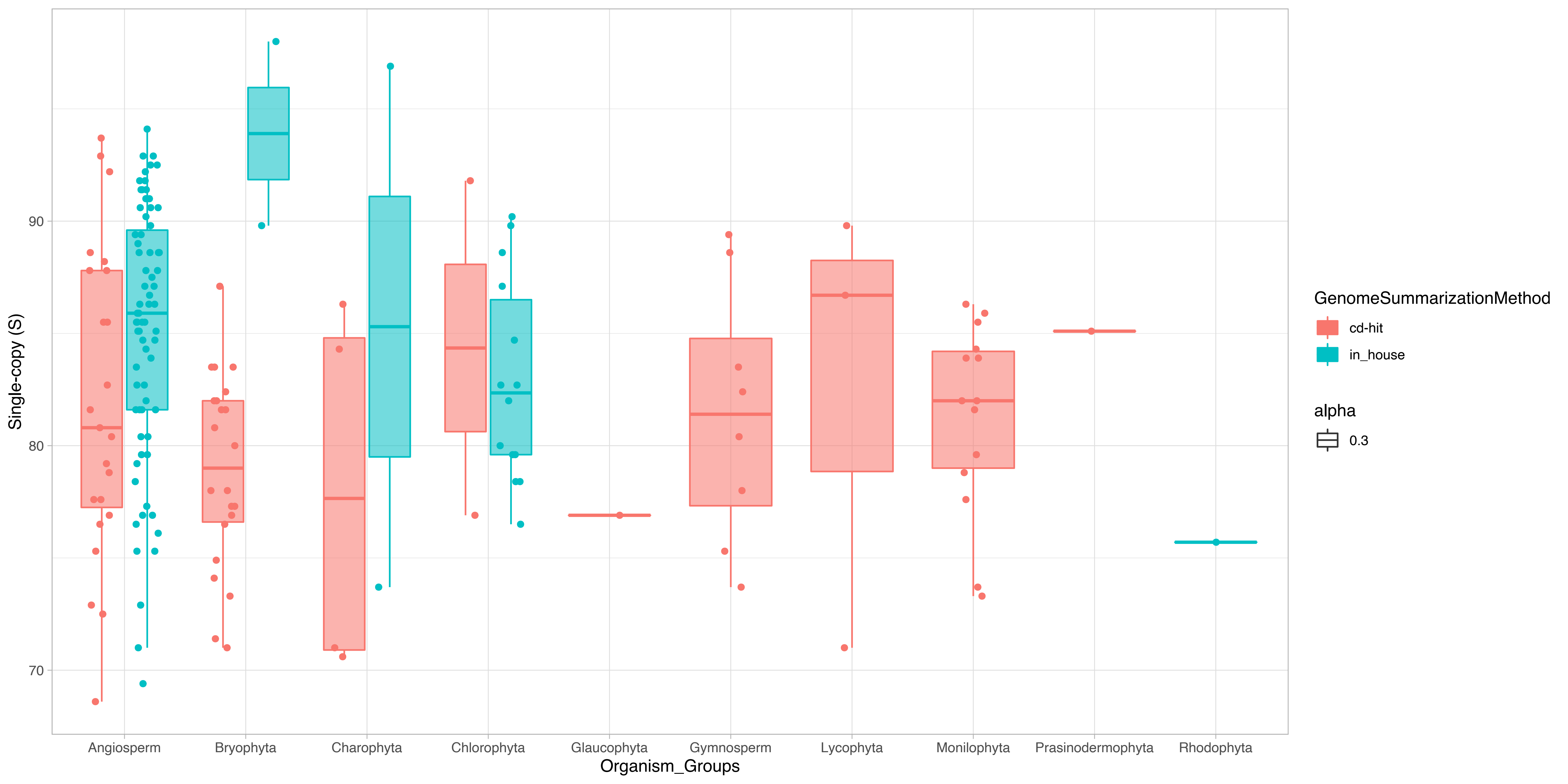

### Fig. S5

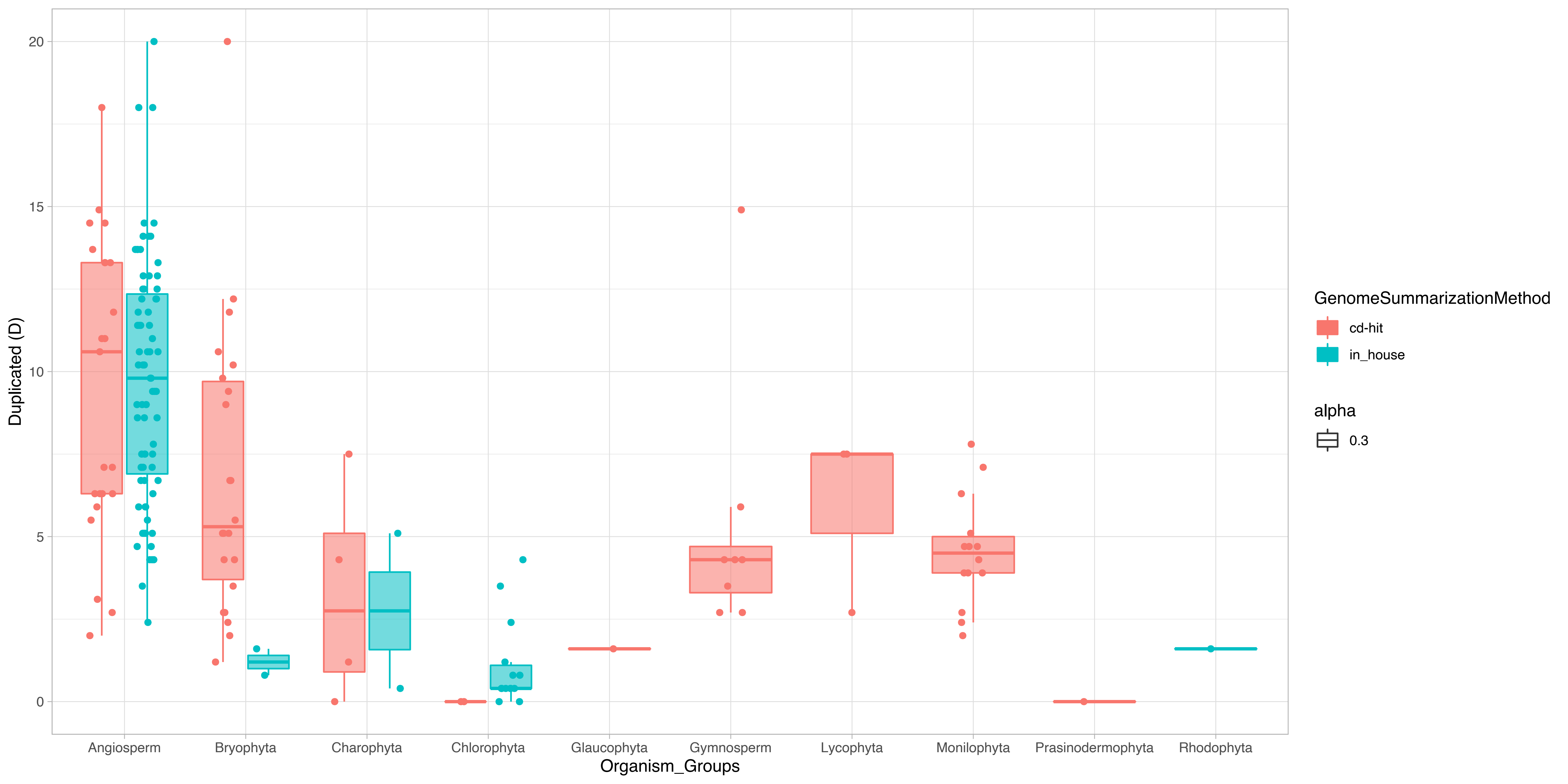

### Fig. S6

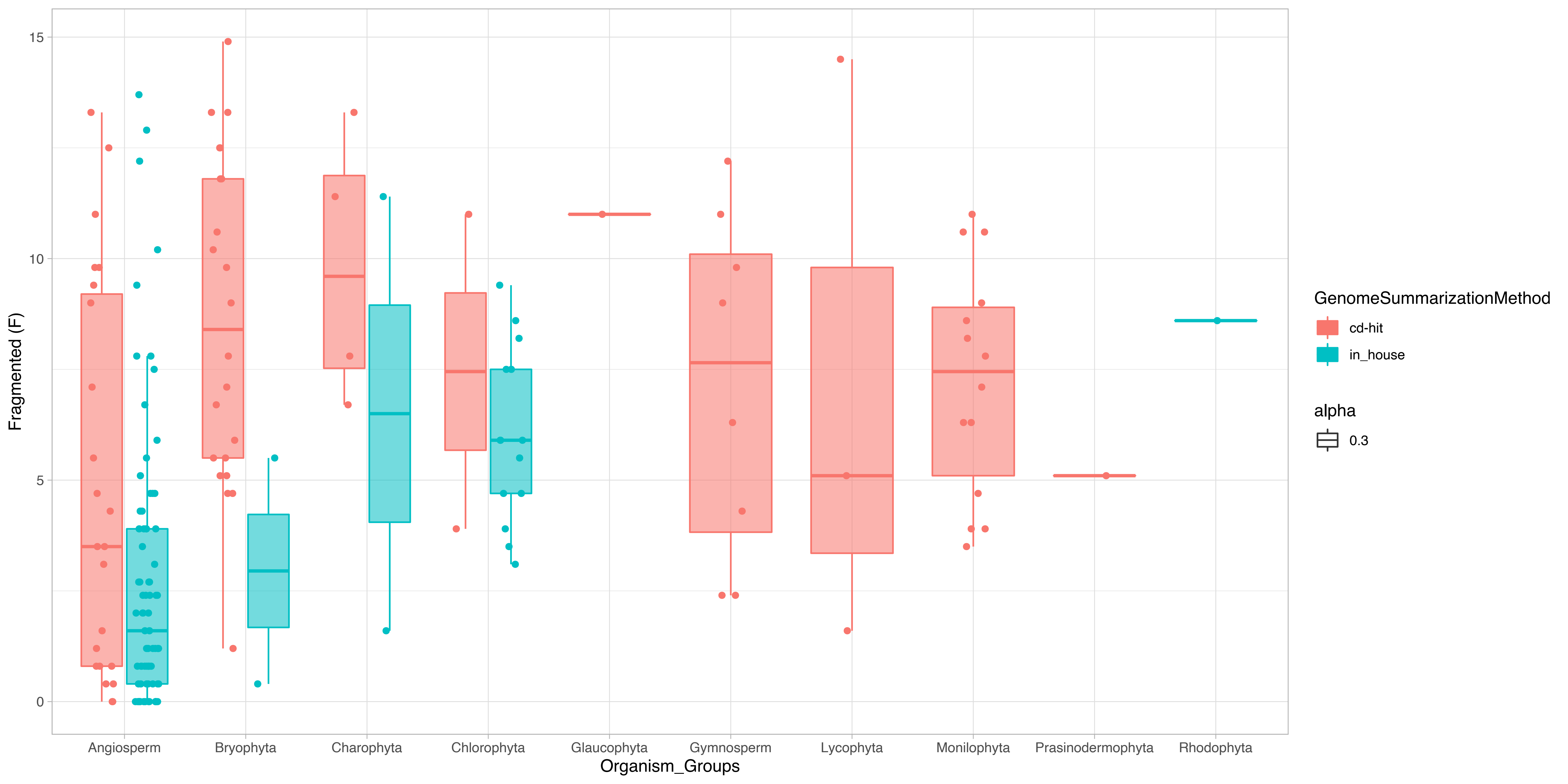

### Fig. S7

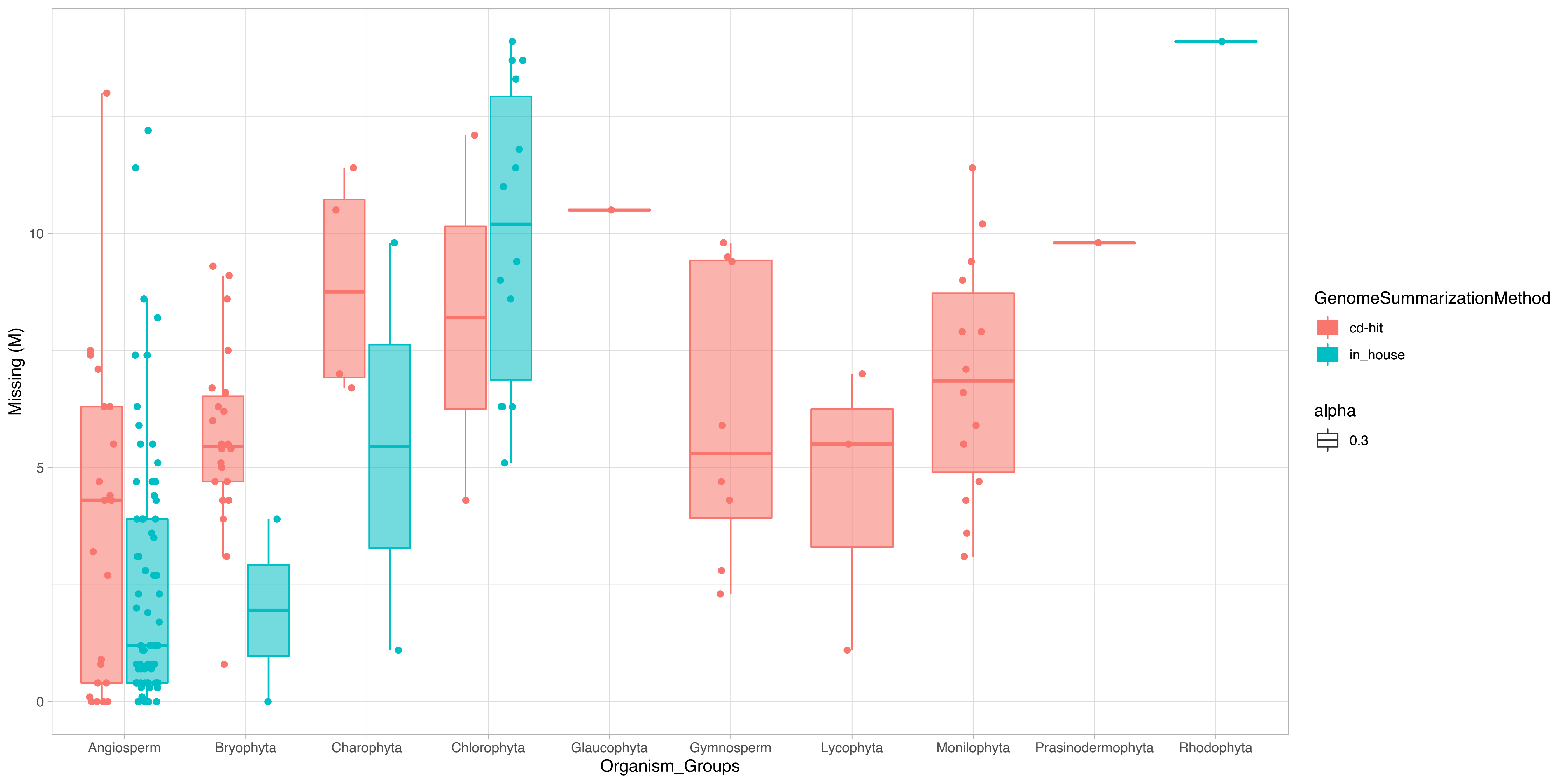
